# Topographical polarity reveals continuous EEG microstate transitions and electric field direction in healthy aging

**DOI:** 10.1101/2024.03.24.586427

**Authors:** Shiho Kashihara, Tomohisa Asai, Hiroshi Imamizu

## Abstract

EEG microstate sequences, representing whole-brain spatial potential distribution patterns of the EEG, offer valuable insights for capturing spatiotemporally continuous and fluctuating neural dynamics with high temporal resolution through appropriate discretization. Recent studies suggest that EEG microstate transitions are gradual and continuous phenomena, contrary to the classical view of binary transitions. This study aimed to update conventional microstate analysis to reflect continuous EEG dynamics and examine differences in age-related electrophysiological state transitions. We considered the relative positions of EEG microstates on the neural manifold and their topographical polarity. Transition probability results showed fewer transitions on the microstate D-C-E axis in older adults. In contrast, transitions among microstates A, D, and B increased in the older group and were mainly observed within polarity. Furthermore, the 100 microstate transitions, which are variations of the shortest transitions between 10 microstates, could be reduced to 8 principal components based on the co-occurrence of each transition, including hubs C and E, planar transitions through msA/B and D, and unidirectional transition components. Several transition components were potentially significant predictors of age group. These features were nearly replicated in independent data, indicating their robustness in characterizing age-related electrophysiological spatiotemporal dynamics.

## 1. Introduction

The resting brain is recognized for its structured activity, linked to various cognitive functions, mental states, and psychological characteristics. This activity demonstrates that the resting brain is neither inactive nor merely random in the absence of stimuli (Greicius, 2008; Gusnard et al., 2001; Miall & Robertson, 2006; Northoff et al., 2010; Stevens & Spreng, 2014). One of the most significant examples of the many aspects related to the resting brain is aging (Andrews-Hanna et al., 2007; Ferreira & Busatto, 2013; Hausman et al., 2020; Varangis et al., 2019). This paper focuses on discretizing the spatiotemporal dynamics of resting electroencephalography (EEG) using a newly developed methodology, with a focus on the effects of aging.

Accumulated studies have revealed that aging is accompanied by various changes in the brain (Aron et al., 2022; Cleeland et al., 2019; Li et al., 2001; Oschwald et al., 2019; Samson & Barnes, 2013). Not only age-related changes in brain structure (Good et al., 2001; Lebel et al., 2010; Oschwald et al., 2019; Resnick et al., 2003) and resting-state functional magnetic resonance imaging (fMRI) functional connectivity (Ferreira & Busatto, 2013; Ferreira et al., 2016; Geerligs et al., 2015; Sala-Llonch et al., 2015), but also age-related differences in the brain are manifested in the electrophysiological neural aspect. For example, research has repeatedly shown that the alpha band frequency activity in the resting-state EEG becomes slow, and its power decreases with age (Babiloni et al., 2006; Chiang et al., 2011; Ishii et al., 2017; Klimesch, 1999; Merkin et al., 2023; Scally et al., 2018; Tröndle et al., 2023).

Despite the growing body of research in cognitive neuroscience related to aging, the spatiotemporal dynamics of whole-brain states with aging remain unclear. EEG is a powerful tool for capturing temporal neural dynamics with good temporal resolution. However, most EEG studies have focused on time- and/or frequency-domain activity in local spatial areas and did not consider whole-brain dynamics. One method for capturing the global spatiotemporal dynamics of whole-brain EEG is EEG microstate analysis (Khanna et al., 2015; Michel & Koenig, 2018). EEG microstates are spatial patterns of scalp potential topographies that transition metastably within a time frame of approximately 40–120 milliseconds (Kleinert et al., 2024; Michel & Koenig, 2018). They reflect the large-scale neural dynamics of brain networks predominant in scalp EEG (Zanesco et al., 2020). Traditionally, four classical microstates have been defined: A, B, C, and D (Michel & Koenig, 2018). Additionally, several additional states are sometimes referred to: C’, E, F, or G (Custo et al., 2014, 2017; Tarailis et al., 2021, 2024). The naming of state labels has diverged across studies due to independent definitions by researchers or naming based on visual similarity or similarity indices with labels reported in previous studies. However, some integration has been attempted recently (Tarailis et al., 2024). The relationship between microstates and function and state of mind continues to interest researchers (Britz et al., 2010; Khanna et al., 2015; Milz et al., 2016; Musso et al., 2010; Panda et al., 2016; Seitzman et al., 2017; Tarailis et al., 2024; Yuan et al., 2012).

Studies examining the relationship between EEG microstates and aging suggest that aging alters specific states, including microstates C, D, and E. In early studies, Koenig et al. (2002) and Tomescu et al. (2018) focused on developmental changes in microstates. They preliminarily described that the state duration of microstate C and the transition probability between microstates C and D decreased with age. It is worth noting that microstate D in Tomescu et al. (2018) appeared to be somewhat posterior-active and thus closer to microstate E, according to the relabeling procedure of Tarailis et al. (2024). Furthermore, a recent study by Zanesco et al. (2020) examined age differences in EEG microstate dynamics using a large open dataset. They found that older adults had larger global explained variances (GEVs) for microstates A and B, smaller GEVs for microstates C and E, and longer average durations and less occurrence in all microstate classes than younger adults. Furthermore, they also suggested that transitions from one state to microstates A and B were more frequent in older adults, while transitions from one state to microstate C and from C to E were fewer in older adults (Zanesco et al., 2020). Additionally, Jabès et al. (2021) revealed that older people have lower GEVs, occurrences, and transitions from one state to microstates C and C’ (noting that their C’ can be relabeled to E; cf., Tarailis et al., 2024).

While findings on age-related microstates are emerging, conventional microstate analysis methodologies have several limitations in providing an adequate discretized representation of continuous EEG spatiotemporal dynamics. The first problem lies in that the labeling of microstates ignores topographical polarity. Conventional microstate analysis has ignored topographical polarity to focus on the electric field pattern in resting-state EEG analysis (Michel & Koenig, 2018; Murray et al., 2008). This technique of ignoring polarity aids in suppressing the impact of unstable and noisy electric fields associated with the inversion of neuronal generators providing EEG oscillations and identifying the persistence of stable states irrespective of the dominant frequency of EEG oscillation (Michel & Koenig, 2018). However, it may not be ideal for characterizing the essentially continuous EEG dynamics in the representation of broad-band spatiotemporal dynamics as a linear sum of activity over multiple frequency bands. Changing topography can indicate changes in global network activity since differences in the spatial representation of the potential map imply differently distributed neuronal generators’ activation (Khanna et al., 2015; Michel & Koenig, 2018). However, the same topography does not imply the same neuronal generator activation. Such an interpretive problem also applies to polarity-reversed topographical patterns. In other words, reversing neuronal generators in the brain may produce polarity-reversal topographies, but this does not mean that all polarity-reversal topographies originate from the same neural generators as the immediately preceding topography. The polarity of the EEG depends on the structural constraints of the brain (e.g., the orientation of the pyramidal cells), instrumental constraints (e.g., electrode orientation and reference position), and changes in E/I balance as the balance between excitatory and inhibitory neurotransmission, as well as the EEG oscillation (Kirschstein & Köhling, 2009). Hence, it is reasonable to assume that polarity in EEG topography has necessary information reflecting the composition of the whole-brain electric field potentials. Therefore, fitting microstate labels to the data in consideration of topographical polarity can aid in understanding the characteristics of EEG spatiotemporal dynamics.

The second problem is that previous microstate studies have relied primarily on static indices. While several quantification indices for EEG microstates exist, the most commonly used measures, such as duration (i.e., how many seconds a state lasts on average) and occurrence (i.e., how often a state occurs on average per second), are static. They are summaries of state characteristics, such as how often or how long a state appears during a specific period, which do not reflect the spatiotemporal continuity of the EEG dynamics. Moreover, these static indices are sensitive to the analysis software and parameters (e.g., temporal smoothing); thus, they can be unstable. Popov et al. (2023) examined the test-retest reliability of various resting-state EEG indices in younger and older participants. They highlighted the low test-retest reliability of EEG microstate measures, particularly among older individuals. While the stability and reliability of EEG microstate indices remain subjects of debate (Antonova et al., 2022; Khanna et al., 2014; Kleinert et al., 2024; Liu et al., 2020; Popov et al., 2023; Zanesco et al., 2020), indices that focus on temporal transitions between states, such as state transition matrices, might be informative, especially for describing continuous trajectories.

Research findings suggest that EEG microstates and their inter-label transitions appear to be continuous rather than discrete (Mishra et al., 2020; Shaw et al., 2019). Therefore, microstate label sequences can be an effective continuative discretization method for characterizing the originally continuous spatiotemporal EEG dynamics. Asai et al. (2023) have shown that the dynamics of continuous EEG topographies can be visualized as continuous trajectories on spherical neural manifolds defined based on dissimilarities between topographies (see Figure S1).. In their methods, neural states are distributed on a closed spherical surface based on the dissimilarity between states, and EEG microstates are represented as attractors in the EEG state space (Férat et al., 2022; Milz et al., 2017). In this context, describing EEG time series as transitions between microstate labels (e.g., AACCCDDDBCC…) can be compared to dividing a closed neural manifold (“the earth;” spherical surface) into several areas (“countries;” arbitrary boundary) and then describing the entry and leave history between countries (“immigration control”). Thus, examining EEG microstate transition offers a suitable approach for discretizing the continuous spatiotemporal dynamics of EEG. Importantly, since each neural state has a topographical polarity, labeling that ignores this polarity leads to the “jumping” of states (Mishra et al., 2020), which cannot account for the continuous trajectories on the spherical manifold. In other words, the two problems of considering topographical polarity and focusing on state transitions are interrelated; therefore, labeling EEG microstates by considering topographical polarity is necessary to properly quantify spatiotemporally continuous dynamics by focusing on state transitions.

In this study, we extended the EEG microstate approach by considering the topographical polarity, especially from the viewpoint of state transitions. We examined the differences in EEG discrete state transitions between younger and older participants. To begin, utilizing a large open dataset (Babayan et al., 2019), we explored the characteristics of the transition matrix between the younger and older adults when topographical polarity was considered, checking for consistency with previous findings of fewer transitions to microstates C and E in the older adults (Study 1). In addition to examining individual transition probabilities between microstates, we attempted to represent coarse-grained EEG dynamics in terms of units of “transition components,” which were groups of similar transitions. This enabled us to identify consistent effects regardless of the unit of measurement scale when examining the relationship between neurodynamic and psychodynamic aspects such as aging. Furthermore, utilizing independent data, we verified whether the transition characteristics between younger and older adults identified in Study 1 could be reproduced at each index scale (Study 2). In this paper, we used the term “positive (+)” topographical polarity, defined as a pattern in which the overall potential of the frontal electrode population is positive and the occipital is negative, and the opposite is termed “negative (-)” topographical polarity (cf., Asai et al., 2022).

## 2. Study 1: Exploratory analysis of topographical polarity considerations using an open dataset

Study 1 utilized the LEMON dataset (Babayan et al., 2019) to compare the transition matrices of microstate label sequences between age groups by considering topographical polarity. This large open dataset has been used in previous studies on EEG microstates (Zanesco et al., 2020). Despite some differences from the study by Zanesco et al. (2020), such as data exclusion criteria and the parameters of the preprocessing and microstate analysis, we expected to replicate their findings that transitions to microstates C and E reduced and transitions to microstates A and B increased in older adults. Moreover, we attempted to extend Zanesco et al. (2020) and other previous studies by considering topographical polarity. By describing microstate sequences considering topographical polarity, we could quantify the transitions between EEG discrete categories while preserving the continuous spatiotemporal dynamics, as expressed as trajectories on neural manifolds (Asai et al., 2023). Previous studies have suggested that the temporal dynamics of EEG are slower and more unitary in older people (e.g., lower peak frequency of oscillation [Ishii et al., 2017; Merkin et al., 2023; Scally et al., 2018] and lower signal variability, i.e., higher regularity in a fundamental dynamic system [Deery et al., 2023; Lau et al., 2022; Ma et al., 2021]). Therefore, the extended microstate sequence used in the present study, which enables us to follow more detailed transitions while accounting for topographical polarity, may emphasize differences in spatiotemporal transition dynamics characteristic of younger and older adults. Hence, to determine what differences in transition dynamics exist between younger and older adults, we conducted an exploratory analysis without specific hypotheses regarding the differences in transitions that account for topographical polarity.

### 2-1. Methods

#### 2-1-1. Data description

We accessed an openly available, published dataset called LEMON (Babayan et al., 2019), consisting of 227 adults categorized into two age groups: younger (20–35 years of age; *N* = 153; female, *n* = 45; male, *n* = 108; mean age = 25.1 years, *SD* = 3.1) and older (59–77 years of age: *N* = 74; female, *n* =37; male, *n* = 37; mean age = 67.6 years, *SD* = 4.7) groups. The large-scale study conducted by Babayan et al. (2019) involved questionnaires, interviews, resting-state EEG and fMRI measurements, cognitive tasks, and other assessments over several days. Data were collected by Babayan et al. (2019) as a part of the Max Plank Institute Leipzig Mind-Brain-Body study database; according to Babayan et al. (2019), their study was conducted in accordance with the Declaration of Helsinki and was approved by the ethics committee at the medical faculty of the University of Leipzig (reference number 154/13-ff). The open data they published were documented, curated, and made available for research purposes (Babayan et al., 2019).

Our interest was in resting-state EEG data. The resting-state EEG was recorded for 16 minutes, consisting of eight consecutive blocks of 1 minute each of eye-closed and eye-opened rest in interleaved order. A 62-channel (61 scalp electrodes according to the international standard 10-10 system and one vertical EOG electrode below the right eye) actiCAP and BrainAmp MR plus amplifier (Brain Products GmbH, Gilching, Germany) were used for EEG recording, with a sampling rate of 2500 Hz. More details are available in Babayan et al. (2019). We downloaded openly accessible raw data (*N* = 215), categorized into younger (*N* = 144; female, *n* = 44; male, *n* = 100; estimated mean age = 22.9 years, *SD* = 3.4) and older (*N* = 71; female, *n* = 35; male, *n* = 36; estimated mean age = 65.4 years, *SD* = 4.6) groups. The LEMON dataset only provides participant ages as discretized 5-year bins for anonymity. Therefore, the minimum value of the discretized bin was used to calculate the estimated mean age and *SD* of each age group (cf., Zanesco et al., 2020). It is noteworthy that for the 55–60 years age bin, all participants were considered 59 years old because the age range for older participants was 59–77 years (Babayan et al., 2019).

#### 2-1-2. Preprocessing of resting-state EEG data

After preprocessing, data from 25 participants (14 younger and 11 older participants) were excluded for the following reasons: extremely small data sizes of the downloaded data files compared to the others, 10 participants (3 younger and 7 older); lack of marker file, 1 younger participant; something wrong with triggers, 2 younger participants; and positive drug tests, 12 participants (8 younger and 4 older). Preprocessing was applied for the 16-minute continuous resting-state EEG data, which comprised a mixture of eye-closed and eye-opened blocks. Our preprocessing procedure is described below. All analyses were conducted using EEGLAB v2019.1 (Delorme & Makeig, 2004) under MATLAB R2019a (Mathworks Inc., Natick, MA, USA).

First, we downsampled data to 250 Hz and applied a Hamming windowed sinc finite impulse response (FIR) filter from 2 to 20 Hz using the EEGLAB function “*pop_eegfiltnew*” of the EEGLAB plugin “*firfilt* (v2.4)”. We removed the vertical EOG channel and then conducted bad channel identification. We employed the artifact subspace reconstruction method (ASR: Mullen et al., 2015) with a threshold of 20 *SD* (Chang et al., 2018) using the EEGLAB function “*clean_artifact*” of the EEGLAB plugin “*clean_rawdata* (v2.3).” The criteria for detecting bad channels were as follows: flat signal longer than 5 seconds, line noise greater than 4 *SD* from the entire signal of that channel, and poor correlation with adjacent channels by less than 85%. These bad channels were removed and interpolated using the EEGLAB function “*pop_interp*.” Under these criteria, an average of 2.62 (*SD* = 3.05, range = 0–16 channels) were detected as bad channels. In addition, ASR results showed that an average of 91.92% (*SD* = 6.40, range = 66.37–99.91%) of the original data were preserved, and the rest were reconstructed as containing non-stationary artifacts.

Subsequent to re-referencing the data to the average potential, adaptive mixture independent component analysis (Hsu et al., 2018; Palmer et al., 2011) was applied using the EEGLAB plugin “*AMICA* (v1.5.1).” Dipole estimation was also conducted using the EEGLAB functions “*dipfit* (v3.3)” and “*fitTwoDipoles* (v0.01).” We then calculated the probability of each of the seven source categories for each independent component (IC) using the ICLabel toolbox (Pion-Tonachini et al., 2019) within the EEGLAB plugin “*ICLabel* (v1.3).” Only ICs that met all three of the following criteria were retained as ICs reflecting the signal and returned to the electrode space for further analysis: ICs labeled “brain” by *ICLabel*, ICs with residual variance lower than 15% (Artoni et al., 2014), and ICs for which dipole was estimated in the brain. A mean of 28.40 ICs (*SD* = 6.71, range = 7–46 ICs) met these criteria.

After data cleaning, the data were divided into eye-opened and eye-closed blocks (one minute each, a total of eight blocks per condition). The eight eye-opened and eye-closed blocks were concatenated and treated as eight-minute resting data. Finally, a total of 190 datasets, including 130 in the younger group (female, *n* = 37; male, *n* = 93; estimated mean age = 22.8 years, *SD* = 3.3) and 60 in the older group (female, *n* = 30; male, *n* = 30; estimated mean age = 65.2 years, *SD* = 4.5) were used in subsequent analyses.

#### 2-1-3. EEG microstate analysis

We conducted EEG microstate analysis using some of the functions of the EEGLAB plugin “*MST1.0*” (Poulsen et al., 2018). The microstate analysis consisted of two phases: a clustering phase to find dominant configuration maps using eye-closed resting data from all 190 participants and a template-matching phase in which the obtained templates were applied to the data of each participant to yield microstate label sequences. It should be noted that temporal smoothing was not applied during EEG microstate analysis in Studies 1 and 2, as our focus was on describing the natural state of continuous spatiotemporal dynamics.

We first calculated the global field power (GFP) time series to extract data for the clustering process. We identified the topography of the local GFP maxima (i.e., peak) for each participant’s EEG time series. GFP is a reference-independent measure of response strength, calculated as the standard deviation of all electrodes at a given time (Murray et al., 2008; Poulsen et al., 2018). The GFP peak point is known to maximize the signal-to-noise ratio of the topography and provide an optimal representation of quasi-stable spatial topographies (Khanna et al., 2015; Mishra et al., 2020; Zanesco, 2020). Of the topographies at the GFP peak points for each participant, 192 were randomly selected per participant for the younger group and 417 topographies for the older group, with a constraint of a 10 ms minimum peak interval. This weighting aimed to correct for the effect of the different number of participants in the younger and older age groups on the microstate templates. It was set up to result in approximately 50,000 data points for clustering, with equal contributions from both age groups (i.e., [50,000 points/2 groups]/130 younger participants ≈ 192 points, and [50,000 points/2 groups]/60 older participants ≈ 417 points).

Subsequently, non-hierarchical clustering was performed on the extracted GFP peak topographies (49,980 topographies in total) using the modified *k*-means method (Mishra et al., 2020; Pascual-Marqui et al., 1995) (k = 2:8, 50 random initializations of algorithm, and 1,000 maximum iterations) using the “*pop_micro_segment*” function in the EEGLAB plugin “*MST1.0*” (Poulsen et al., 2018). After that, we determined the optimal number of clusters based on the cross-validation (CV) criterion (Murray et al., 2008; Pascual-Marqui et al., 1995). The resulting microstate template maps corresponded to a unit-length basis vector defining a 1-D subspace embedded in a *k*-dimensional space (Mishra et al., 2020). We aimed to incorporate topographical polarity into the template-matching process for the EEG time series data maps. To achieve this, we opted to use twice the optimal number of *k* maps obtained using the modified *k*-means algorithm. This encompassed both the original maps and their inverted counterparts, resulting in a total of k*2 microstate templates used in this study.

Next, the template maps were matched to the original EEG recordings for each participant to yield a time series sequence of microstates using the “*pop_micro_fit*” function in the EEGLAB plugin “*MST1.0*” (Poulsen et al., 2018). For all sampling points comprising each participant’s eye-closed resting-state EEG time series, the spatial correlations between the template maps and the topography map of each sampling point were calculated. Subsequently, the topographical map of each point was classified into template labels that showed the highest spatial correlation (“winner-take-all”). Importantly, when considering topographical polarity, the assignment of template labels involved referencing a template map that included both the original map and its polarity-reversed counterpart. In addition, no temporal smoothing was applied.

The obtained microstate label sequences were divided into transitions between two consecutive states using sliding windows of two-time points with 50% overlap. The number of transitions was counted for a total of 100 transitions: the combination of [optimal number of *k*×2] previous states (“from” state) × [optimal number of *k*×2] following states (“to” state).

In terms of comparing age differences for each transition pair, we defined a transition probability matrix as follows: the transition probability matrix was calculated by averaging the transition probability matrix of each individual obtained by first counting the number of microstate label transitions between two consecutive time points and then dividing by the total number of state transitions excluding self-recurrences. Various definitions exist for transition matrices (e.g., whether to use the number of occurrences or the probability, whether to calculate the overall probability of 1 or the probability from a certain “from” state, and whether to allow self-recurrence to the same label). Here we employed a transition probability with all transitions except self-recurrences set to 1, to focus on the switching of a certain electric field pattern, i.e., the ease or difficulty of switching in the neural network. Readers may refer to the supplementary materials for results when other transition matrix definitions were applied.

#### 2-1-4. Statistical analysis

In Study 1, we conducted an exploratory analysis of differences in EEG transition dynamics between younger and older adults. To explore the different characteristic transitions between older and younger groups, we calculated the effect size of the *d* family on the difference in transition probabilities for each transition in the two age groups (standardized mean difference obtained by dividing the difference in mean transition probability by the standard deviation of the two groups pooled; Hedge’s *g*), as described by Zanesco et al. (2020). For differences in transition probabilities between age groups, we treated effect sizes greater than or equal to the required effect size determined by power analysis as “meaningful differences” (Zanesco et al., 2020). We conducted power analysis using G*Power 3.1.9.7 to evaluate the sensitivity in the difference between the two age groups. Power analysis revealed that an effect size ≳ 0.44 could be detected with an alpha error probability of 0.05, a power (1-β) of 0.8, and sample sizes of 60 and 130 for the older and younger age groups, respectively.

In addition to examining age differences in the transition probability of each transition pair, we conducted a principal component analysis (PCA) on the number of transitions for all 100 transitions for all participants to characterize the microstate transitions. Applying scaling and centering to the number of counted transitions and then PCA, the inter-participant correlation matrix for the number of transitions for all 100 transition pairs was transformed into an uncorrelated composite variable. We then applied Varimax rotation after PCA to examine the dimensional structure along the original data structure and improve the data’s interpretability. PCA was performed in RStudio version 2021.09.0 and R version 4.3.1 (R Core Team, 2023) using the R function “*prcomp*.” Subsequently, the relationship between the age groups and the components of transitions was examined in both directions using logistic regression and multivariate *t*-tests (Hotelling’s *T*^2^ test), utilizing the R functions “*glm*” and “*HotellingsT2Test.*” We used the R function “*effectsize*” to calculate the Hedge’s *g* for post hoc Welch’s *t*-test. We applied Holm’s method to adjust the significance level in multiple comparisons. Adj.*p* in the results section represents the adjusted *p*-value, obtained by dividing the conventional significance level of 5% by the number of comparisons for convenience.

### 2-2. Results

#### 2-2-1. EEG microstate templates

After modified *k*-means clustering, five maps were extracted from all participants’ eye-closed data, in line with the approach used by Zanesco et al. (2020). These five maps explained 78.21% of the GEV. This value was comparable to those observed in other studies: a previous review showed that microstate studies employing four cluster maps explained approximately 65–84% of the GEV (Michel & Koenig, 2018). We labeled them as microstates A, B, C, D, and E by careful visual inspection based on the labels unified in Tarailis et al. (2024) (Figure 1a). The spatial correlations between our templates and those of a previous study (Asai et al., 2022), which employed the four template maps A, B, C, and D, were sufficiently high (for microstates A–D, *r*s > .88; Figure 1b). Additionally, our microstate E had the highest correlation coefficient with microstate C among the four maps used by Asai et al. (2022) (*r* =.73), consistent with the findings of previous studies that microstates C and E had not previously been distinguished (e.g., Tarailis et al., 2021).

**Figure 1.**
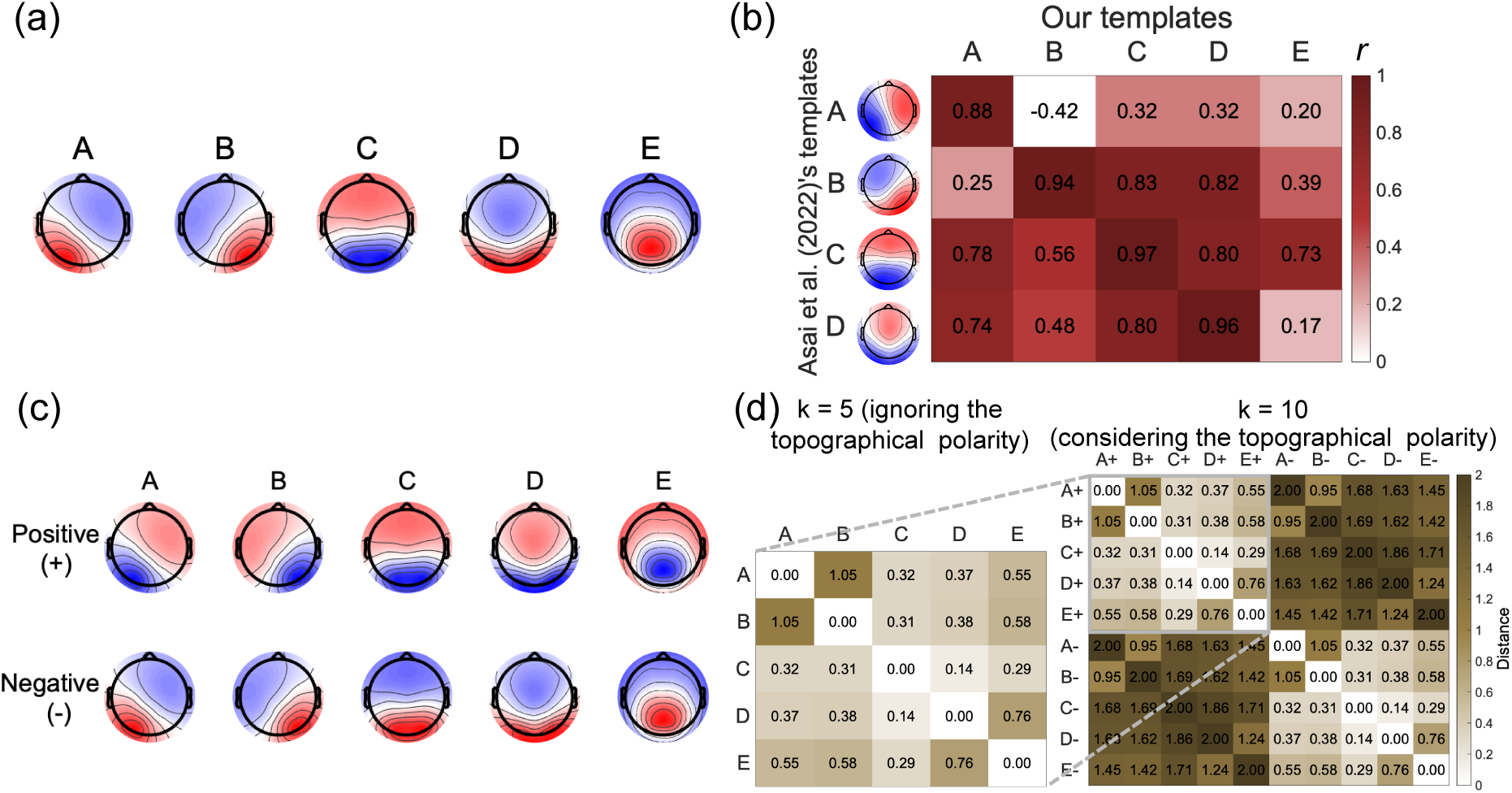
EEG microstate templates. (a) Spatial distribution of clustering maps extracted from the eye-closed resting-state data of all 190 participants in the LEMON dataset. We selected k = 5 as the optimal number of k according to the CV criteria. Labels A–E were assigned based on similarities with templates identified in Asai et al. (2022) and through visual inspection based on Tarailis et al. (2024), who unified microstate labels. (b) Example of spatial correlation between our templates and those of other studies. We compared our templates with those of Asai et al. (2022). The spatial correlation coefficients were computed using a vectorized 67*67 interpolated scalp map data matrix obtained using the EEGLAB function “topoplot” to absorb differences in the number of electrodes. In extracting microstate templates as basis vectors at this point, we ignored topographical polarity, i.e., we only considered spatial configuration. For instance, the topographical polarity of microstate A in Asai et al. (2022) was reversed from our microstate A, so we aligned both maps to positive topographical polarity and then calculated their spatial correlations for convenience. (c) We used the polarity-considered template maps. Each map extracted by the k-means method was inverted, and a total of ten maps were used as our microstate templates in this study. (d) The dissimilarity matrix between template maps was defined by the spatial correlation among the five (left) or ten (right) template maps. By subtracting the correlation coefficients from 1, we could transform a correlation matrix into a dissimilarity matrix (one of the definitions of distance) between the topographies. These dissimilarities range from 0 to 2; values closer to 0 indicate closer distances between two topographies (0 means the two are the same), while values closer to 2 indicate greater differences (2 means that the two have polarity-reversed patterns). This matrix was used to determine the positional relationship of each template map when drawing directed graphs.

Next, to perform template-matching considering the topographical polarity of the maps comprising the EEG time series data, after identifying the *k* (*k* = 5) dimensional space, we decided to consider the positive and negative directions of each dimensional axis (i.e., topographical polarity). Microstates obtained by modified *k-*means clustering can be treated as basis vectors of a subspace of the EEG channel space, i.e., all topographical states on that basis vector show the same electric field pattern (Mishra et al., 2020). We tried to extend this by treating the EEG microstate basis vector as an “oriented” vector that considers orientation. A polarity-reversed map was created and added for each map obtained by the modified *k*-means algorithm. Thus, ten maps were created that were used as microstate templates for polarity-considering matching in this study (Figure 1c).

Furthermore, the positional relationship of each template map was identified based on the distance between maps. Specifically, the positional relationship between maps was determined by compressing the distances defined by the spatial correlation coefficients between topographies (Figure 1d; i.e., dissimilarity matrix between topographies) in three dimensions using the classical multidimensional scaling (MDS) method (Asai et al., 2023; Koenig et al., 2024). We confirmed that when the topographical polarity was considered, the EEG topographies containing EEG microstate templates, which were centroids of states in specific categories, were arranged to cover the surface of the sphere (Figures 2a and 2b). The relative positions of the 10 EEG microstate templates were revealed (Figure 2c).

**Figure 2.**
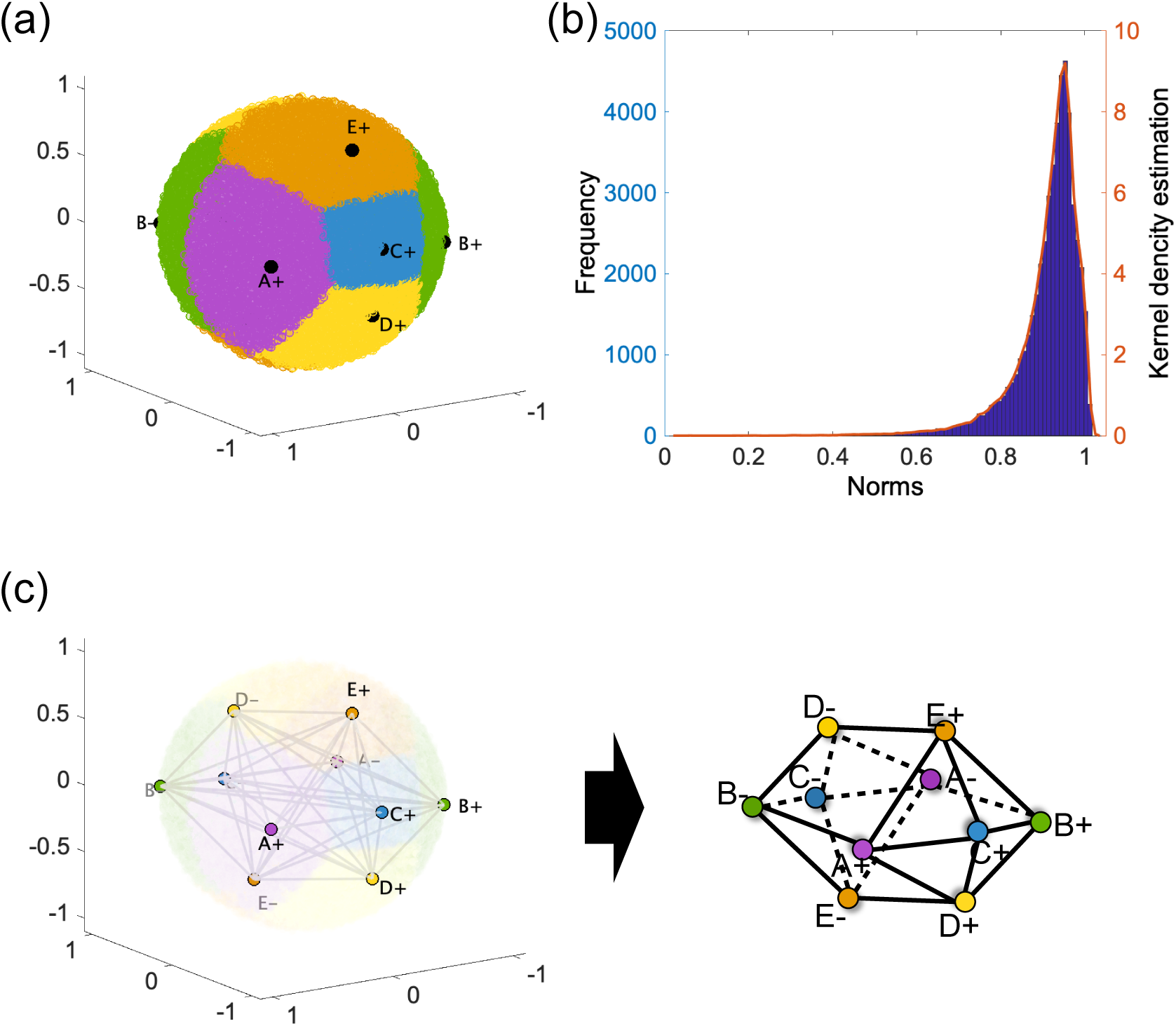
Example of neural manifolds with spatial configurations of scalp potentials distributed in state space using the classical multidimensional scaling method. (a) Distribution of topography over neural state space. The method for creating neural manifolds was based on that described by Asai et al. (2023). Each dot represented one state of the GFP peak data (49,980 topographies collected from all 190 participants for template clustering). The topography map was compressed into 3-D information based on spatial dissimilarities between states and then placed on a sphere. The color of the dots indicated the template cluster to which each state belonged (purple for microstate A, green for B, blue for C, yellow for D, and orange for E). A large black circle in each area meant a microstate template. The area of each template cluster comprised one square (microstate C; blue area) and four trapezoids (microstates A, B, D, and E; purple, green, yellow, and orange areas, respectively), which were pasted together around C to form a hemisphere, resembling a soccer ball. In this representation, the EEG microstate templates were represented as attractors on spherical neural manifolds. Since this was a dissimilarity-based distribution, the same labels with different polarities (e.g., A+ and A-) were located at opposite poles of the sphere. (b) Histogram of the vector norm for each state. Since the neural manifold was theoretically a sphere of diameter two and radius one because it was created from dissimilarities based on correlation coefficients (range, 0–2), a norm closer to 1 indicates that the states are aligned on the spherical surface; thus, most of the states were located on the sphere’s surface. (c) Image of a polarity-sensitive 10-state network graph. Transitions between template maps can be written as discrete network graphs while preserving the positional relationships between microstate template maps in the neural state space.

#### 2-2-2. Replication of previous findings when using the topographical polarity-ignored microstate template maps

First, as a confirmation, we checked whether we reproduced the transition features of the older and younger age groups shown in Zanesco et al. (2020) and many other previous microstate studies, with polarity ignored in our analysis. Previous studies have demonstrated that transitions to microstates C, D, and E were less common, and transitions related to microstates A and B were more common in older than in younger adults (Jabès et al., 2021; Tomescu et al., 2018; Zanesco et al., 2020). We performed a conventional microstate analysis applying the five template maps (before adding inversion maps) and calculated the transition probability matrices between the five category maps for the older and younger age groups. The difference between the transition probability matrices for the age groups and their standardized differences were then calculated. These transition probabilities were also represented as directed graphs with each template map as a node and the frequency of transitions between nodes or the magnitude of the difference between age groups as edges.

The five template maps explained an average of 69.78% (*SD* = 5.77) of the GEV of each participant’s EEG topographic time series. The total GEV could also be divided into GEVs for each of the five categorical maps, with microstate A having a mean GEV of 11.37% (*SD* = 4.08); B, 11.86% (*SD* = 4.28); C, 22.52% (*SD* = 10.32); D, 14.82% (*SD* = 6.19),; and E, 9.20% (*SD* = 4.44).

Figure 3 shows the matrix and directed graph of the transition probabilities of each age group (Figures 3a and 3b), difference between each transition pair for both groups (older-younger; Figure 3c), and standardized mean difference between age groups for each transition pair (Figure 3d). In line with previous findings, we observed that transitions to microstate C were less frequent in the older group than in the younger group. In particular, the fewer transitions from microstates C to E, E to D, and D to C in the older group exceeded the effect size detectable in Study 1 (|*d*| = 0.51-0.64).

**Figure 3.**
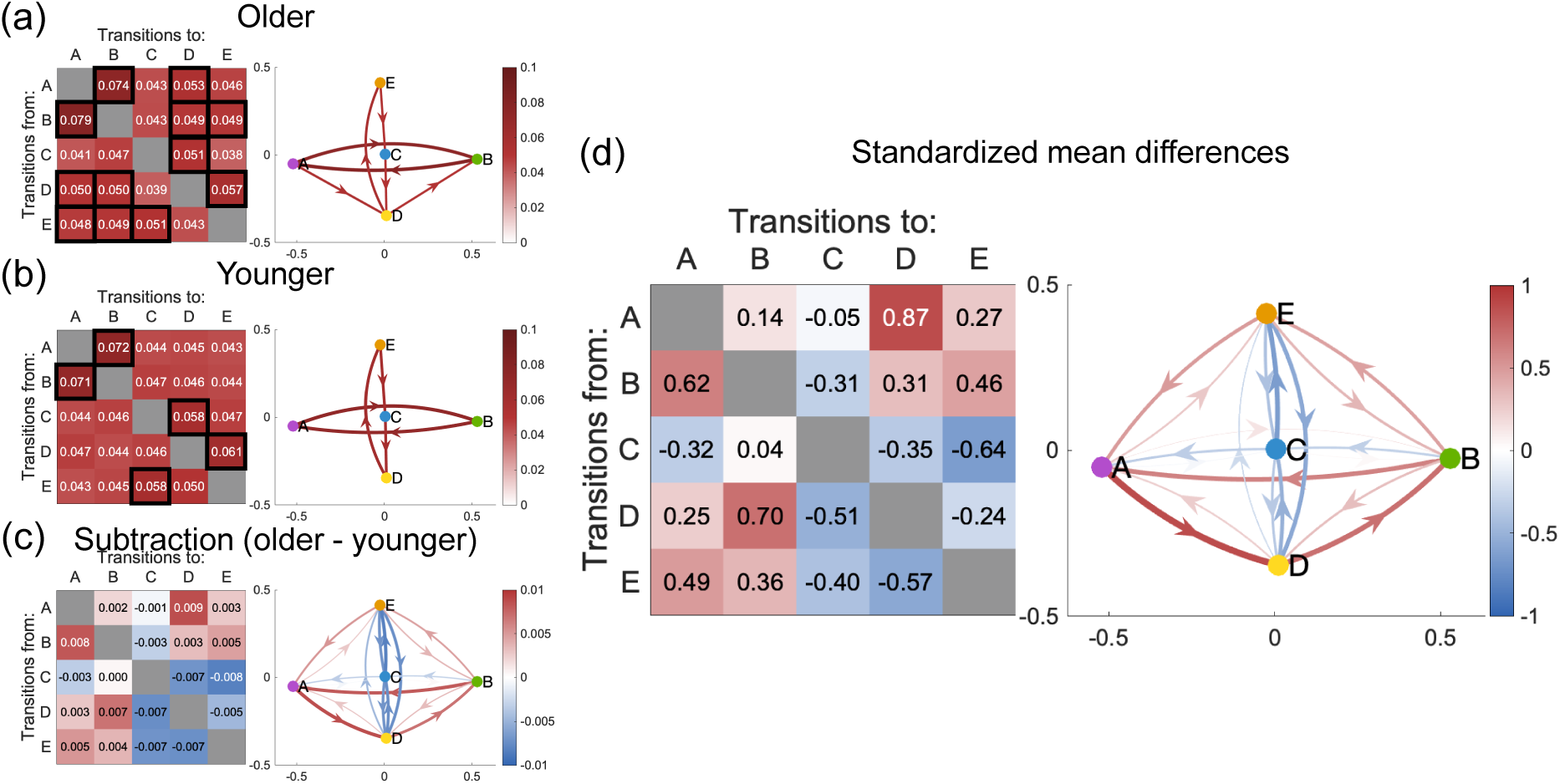
Traditional view of microstate transitions—transition probability matrices and directed graphs of microstate transition pairs ignoring topographical polarity in Study 1. Each transition pair’s average probability of occurrences was shown by placing the “from” states in the row direction and the “to” states in the column direction. Twenty transition pairs were included, except for five self-recurrences on the diagonal. In each directed graph, the nodes were the respective template maps, and the edge color indicated the transition probabilities between nodes or the magnitude of the differences between age groups. The edges of the directed graphs in (a) and (b) were only displayed if the transition probability was greater than 0.05 (= the chance level probability defined as random transitions between all states). Black squares in each transition matrix indicated transitions with valid edges in the directed graph. The distribution of nodes was defined based on the spatial correlations between templates and determined based on the dissimilarity distances compressed by classical MDS (Asai et al., 2023; Koenig et al., 2024). (a) Average transition probabilities for the older and (b) younger groups. (c) Differences in the average transition probabilities for each transition pair for older and younger groups. Red indicates more frequent transitions in the older group, and blue indicates less frequent transitions. (d) Standardized mean differences for comparisons between age groups (Hedge’s g; Zanesco et al. (2020)). The symbol of the effect size g and red-blue colors were used to indicate the direction of the difference, with red indicating more frequent transitions and blue indicating less frequent transitions in the older group.

In addition, transitions related to microstates A and B were conversely more common in the older group than in the younger group. In particular, higher transition probabilities were observed from microstates A to D, D to B, B to A, E to A, and B to E in the older group. These exceeded the effect sizes detectable in Study 1 (|*d*|s = 0.44-0.87).

Interestingly, the directed graphs visualizing the transition matrices also revealed the relative positional relationships among the template maps (Figure 3; see Figure 1d). As depicted in Figure 3, based on the dissimilarities among the five template maps, the topography of the 32-electrode space was compressed, and the five topographies were embeddable in 2D. The relative position of the five template maps was the furthest distance between microstates A and B, followed by microstates E and D. The axes connecting microstates A and B, as well as E and D, were arranged in a cross shape, with distances A-E and B-E, as well as distances A-D and B-D, nearly equal. On the other hand, when comparing distances E-C and C-D, E-C was slightly further apart than C-D. Furthermore, microstate C was located at the intersection of the two axes.

#### 2-2-3. EEG microstate transitions by considering the topographical polarity

Next, we performed an extended microstate analysis, which incorporated template-matching while accounting for topographical polarity. We could obtain ten polarity-considered microstate templates by adding the polarity-reversed topographical maps to the clustering maps (Figure 1c). This allowed us to subdivide transitions into several sub-transitions. For example, the transition from A to B could be subdivided into four transitions: from A+ to B+, from A+ to B-, from A- to B+, and from A- to B-. The ten templates were applied to the data, as in section 2-2-2. This allowed us to obtain results on the transition probability matrices and directed graphs among the ten templates for each age group and the differences between the two groups.

First, the positional relationships among the ten templates were clarified in 3D state space through classical MDS based on spatial correlations among template maps (Asai et al., 2023; Koenig et al., 2024). The positions of five template maps of respective positive or negative polarity were consistent with the results of the five templates without polarity (in section 2-2-2; see Figure 3). Additionally, the templates with reversed polarity (e.g., A+ and A-) were located opposite each other because the dissimilarity value was 2. Ten states were placed on the 3-D compressed abstract space, as shown in Figure 4, and their position was determined (see also Figure 4 and Figure S5 for the animation image).

**Figure 4.**
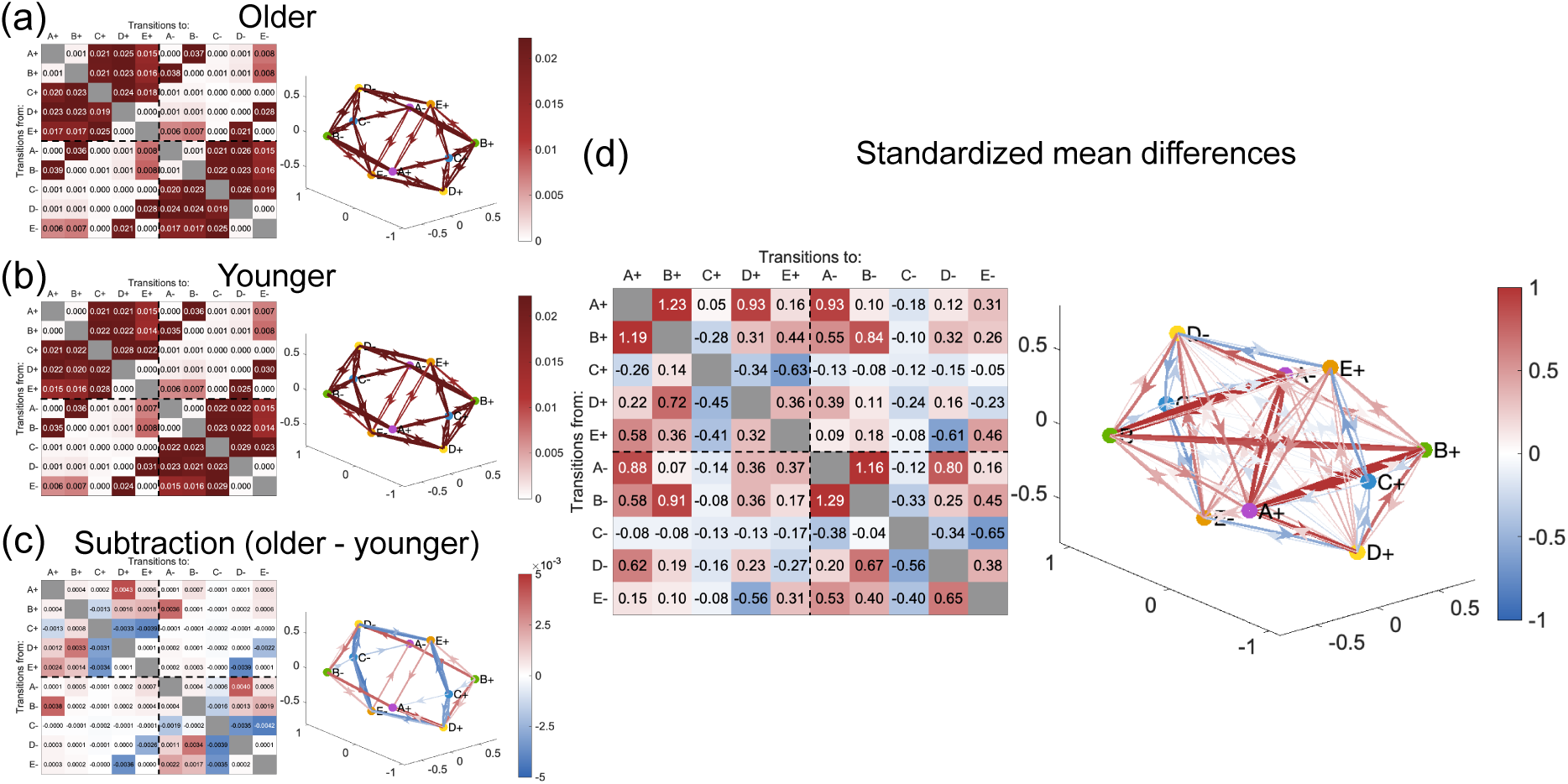
Updated view of microstate transitions—transition probability matrices and directed graphs of microstate transition pairs when considering topographical polarity in Study 1. Each transition pair’s average probability of occurrences was shown by placing the “from” states in the row direction and the “to” states in the column direction. Ninety transition pairs were included, except for ten self-recurrences on the diagonal. In each directed graph, the nodes were the respective template maps, and the edge color indicated the transition probabilities between nodes or the magnitude of the differences between age groups. The edges of the directed graphs were only displayed if the transition probability was greater than 0.011 in (a) and (b) (= the chance level probability defined as random transitions between all states) and greater than 0.0011 in (c) for visualization purposes. The distribution of nodes was defined based on the spatial correlations between templates and determined based on the dissimilarity distances compressed by classical MDS (Asai et al., 2023; Koenig et al., 2024). (a) Average transition probabilities for the older and (b) younger groups. (c) Differences in the average transition probabilities for each transition pair for older and younger groups. Red indicates more frequent transitions in the older group, and blue indicates less frequent transitions. (d) Standardized mean differences for comparisons between age groups (Hedge’s g; Zanesco et al. (2020)). The symbol of the effect size g and red-blue colors were used to indicate the direction of the difference, with red indicating more frequent transitions and blue indicating less frequent transitions in the older group.

Template matching using a total of 10 maps on average explained 69.78% (*SD* = 5.77) of the GEV of the EEG topographic time series for each participant. The GEV of each microstate label was as follows: microstate A+, 5.86% (*SD* = 2.09); B+, 6.15% (*SD* = 2.29); C+, 11.60% (*SD* = 5.48); D+, 7.71% (*SD* = 3.38); E+, 4.74% (*SD* = 2.34); A-, 5.51% (*SD* = 2.01); B-, 5.70% (*SD* = 2.02); C-,10.92% (*SD* = 4.89); D-, 7.11% (*SD* = 2.86); and E-, 4.46% (*SD* = 2.13).

Figure 4 shows the matrix and directed graph of the averaged transition probabilities of the older and younger groups (Figures 4a and 4b), subtraction results between age groups (older-younger; Figure 4c), and standardized mean difference between age groups for each transition pair (Figure 4d). The results suggested that considering topographical polarity during the template-matching process extended the previous findings. When considering the topographical polarity, it was observed that not all of the sub-transitions occurred equally, but there was a bias in the transition probabilities that occurred. For example, transitions from/to C were restricted within the same polarity. Transitions between topographical polarities basically rarely occurred (see the upper right and lower left segments of the transition probability matrices in Figures 4a and 4b). However, some specific transitions (i.e., between A and B, as well as between D and E) were limited to transitions between polarities. For example, transitions from A to B were limited to those from A+ to B- (averaging 3.6% in the younger group and 3.7% in the older group) or from A- to B+ (averaging 3.6% in the younger group and 3.6% in the older group), while transitions from A+ to B+ (averaging 0.0% in the younger group and 0.1% in the older group) or A- to B- (averaging 0.0% in the younger group and 0.1% in the older group) were rarely observed.

Regarding the differences between older and younger adults, it was reproduced that transitions between microstates C, D, and E were less frequent and that transitions associated with microstates A and B were more common in the older group. Such differences were restricted to topographical polarity: for transitions involving microstates C, D, and E, within-polarity transitions (C-D and C-E) and between-polarity transitions (D-E) were observed less frequently in older adults (with effect sizes above detectable thresholds, transition of microstates D+ to C+: |*d*| = 0.45; C+ to E+: |*d*| = 0.63; E+ to D-: |*d*| = 0.61; D- to C-: |*d*| = 0.56; C- to E-: |*d*| = 0.65; E- to D+: |*d*| = 0.56). Plotting these on the neural manifold suggested a circle-shaped decreasing transition route connecting D-C-E sequentially between polarities (Figures 4c and 4d). The C, D, and E-related transitions other than the above, such as between-polarity C-E and C-D transitions or within-polarity D-E transitions, were not significantly reduced among older participants.

Conversely, transitions involving microstates A and B were notably more frequent within the same polarity, primarily characterized by increased transition from A to D (positive: |*d*| = 0.93; negative: |*d*| = 0.80), D to B (positive: |*d*| = 0.72; negative: |*d*| = 0.67), B to E (positive: |*d*| = 0.44; negative: |*d*| = 0.45), and E to A (within positive polarity: |*d*| = 0.58; within negative polarity: |*d*| = 0.53) in older participants (Figures 4c and 4d). Transitions from B+ to A- and B- to A+ were also more common among older participants (|*d*|s = 0.55 and 0.58, respectively).

Furthermore, a characteristic feature of the transitions in older participants was the increased occurrence of transitions that were rare. Specifically, within-polarity transitions between A and B (|*d*|s = 1.16 to 1.29), transition from E- to D- (|*d*| = 0.65), transition from D- to A+ (|*d*| = 0.62), and transitions between opposite same-label states (between A±: |*d*|s = 0.88-0.93; between B±: |*d*|s = 0.84-0.91; from E+ to E-: |*d*| = 0.46), were more common in older than in younger adults.

The number of nodes in the direction between polarities was lower for the 10 templates created in the data-driven manner than in the other axes. Therefore, to ensure accuracy, we further verified the transition matrices using 14 templates, incorporating artificial midpoints between A+ and B+ (designated as Ab), A- and B+ (designated as aB), D+ and E- (designated as De), and D- and E+ (designated as dE). This adjustment was made to ensure that the distances between the transition nodes on the sphere were approximately equal (cf., Figure 2c). As a result, the transition trends observed in the 10 templates were robustly reproduced (Figure 5).

**Figure 5.**
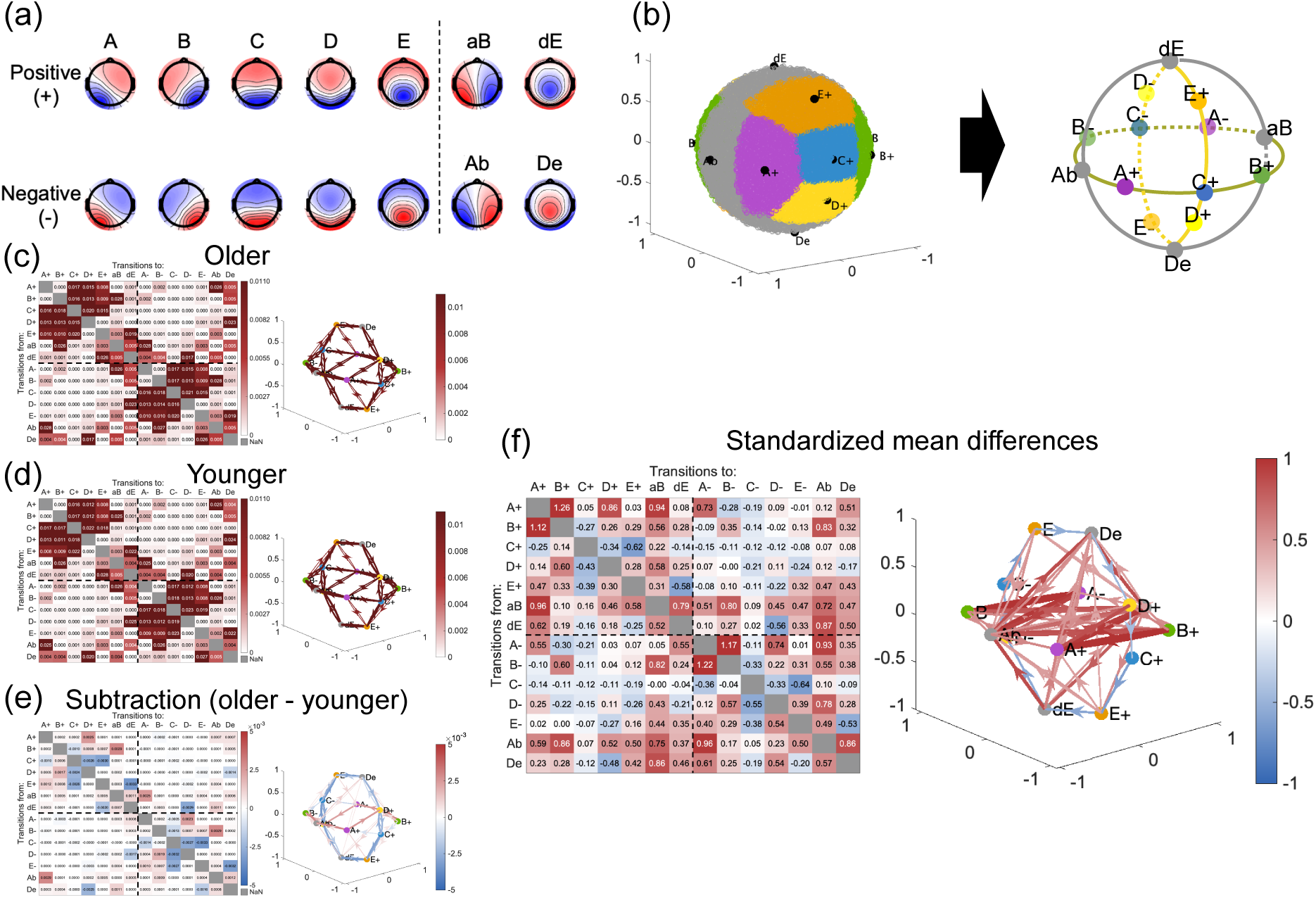
Examples of microstate transition matrices and directed graphs for different node definitions. (a) Example with 14 template maps defined as nodes. (b) Addition of four template maps, including midpoints between A-B and D-E, enabling the arrangement of template maps to evenly divide each of the A-C-B and D-C-E axes. (c) Mean transition probabilities for each of the older and (d) younger groups, (e) differences between the two groups, and (f) standardized mean differences for comparisons between age groups. Details of (c)-(f) are the same as in Figures 4a-d.

To clarify the effect of considering topographical polarity on age group differences, we also compared the transition probabilities for each within- and between-polarity transition from one state to another. We simplified the options of “to” states into two and recalculated the transition probabilities (Figure 6): within- or between-polarity transitions. For within-polarity transitions, we found a significant *T*^2^ effect (*T*^2^ = 42.26, *df* = 10, *p* < .001). Post-hoc analysis revealed that the older group was more likely to transition from microstate A± to other states of the same polarity (A+: *t*(116.93) = 4.86, *p* < .001, adj.*p* < .001, *g* = 0.75; A-: *t*(128.32) = 3.72, *p* < .001, adj.*p* = .002, *g* = 0.55) while less likely to within-polarity transition from microstate C± (C+: *t*(108.36) = -4.29, *p* < .001, adj.*p* < .001, *g* = 0.68; C-: *t*(114.24) = -5.04, *p* < .001, adj.*p* < .001, *g* = 0.79). For inter-polar transitions, we found a significant *T*^2^ effect (*T*^2^ = 23.11, *df* = 10, *p* = .01). Post-hoc analysis revealed that the older group was more likely to between-polarity transition from B± (i.e., mostly from B± to A∓; B+: *t*(95.22) = 3.63, *p* < .001, adj.*p* = .005, *g* = 0.61; B-: *t*(95.40) = 3.59, *p* < .001, adj.*p* = .005, *g* = 0.61) and less likely than the younger group to make an inter-polar transition from E (i.e., mostly from E± to D∓; E+: *t*(164.47) = -2.94, *p* = .004, adj.*p* = .03, *g* = 0.40; E-: *t*(159.07) = -2.62, *p* = .01, adj.*p* = .07, *g* = 0.36).

**Figure 6.**
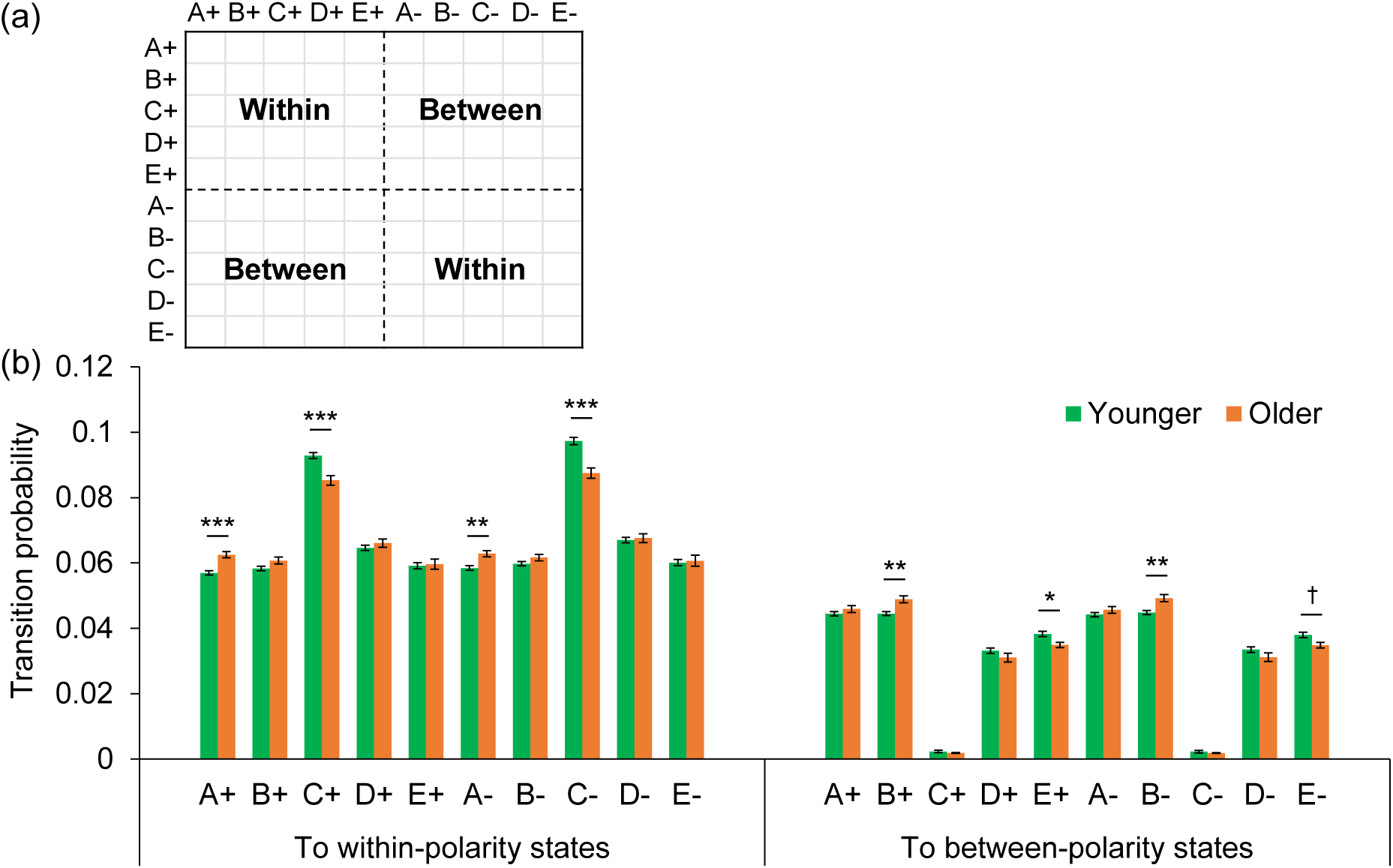
Comparison of transition probabilities from one state to another within and between polarities between age groups in Study 1. (a) Schematic figure of within- and between-polarity transitions in the transition matrix. (b) Within- and between-polarity transition probabilities by age group. The left panel shows the probability of transitioning within polarity from each state (ABCDE±), and the right panel shows the probability of transitioning between polarities from each state. *Note.* ^***^adjusted *p* (adj.*p*) < .001, ^**^adj.*p* < .01, ^*^adj.*p* < .05, ^†^adj.*p* < .10. Error bars indicated the standard errors.

#### 2-2-4. Transition components based on inter-participant correlation of transition probabilities

We performed PCA using the inter-participant correlation matrix for the counts of each transition pair (Figure 7). Based on parallel analysis, eight components were employed (Figure 8a); these eight PCs explained 87.65% of the variance in the data (Figure 8b). After PCA, Varimax rotation was applied to the PCA results to interpret each PC. Each transition pair was grouped as belonging to the component with the highest absolute value of rotated component loadings (see Table S1). The transition pairs contributing to each rotated component after Varimax rotation are shown in Figure 8c. From the list of transition pairs contributing to each rotated component, each component was interpreted and named as follows. Rotated PC1 was named “Hub C” because it was a set of transitions between C (C+/C-) and the opposite ABDE and the self-recurrence of C. Rotated PC2 was named “Hub E” because most of the contributing transition pairs were a set of transitions from/to E (E+/E-). Rotated PC3 was named “E-C-D rotation” because most of the contributing transition pairs were a set of transitions belonging to the one-directional transition routes E+, C+, D+, E-, C-, and D-. Rotated PC4 was named the “A-D plane” because it was a set of transitions between A (A+/A-) and D (D+/D-). Rotated PC5 was named the “B-D plane” because it was a set of transitions between B (B+/B-) and D (D+/D-). Rotated PC6 was named “D-C-E rotation” because it contained a transition route in the opposite direction of rotated PC3. Rotated PC7 was named “A-C-B rotation” because it was a set of transitions belonging to one-directional transition routes A+, C+, B+, A-, C-, and B-. Finally, rotated PC8 was named “B-C-A rotation” because it was a set of transition routes in the opposite direction of rotated PC7.

**Figure 7.**
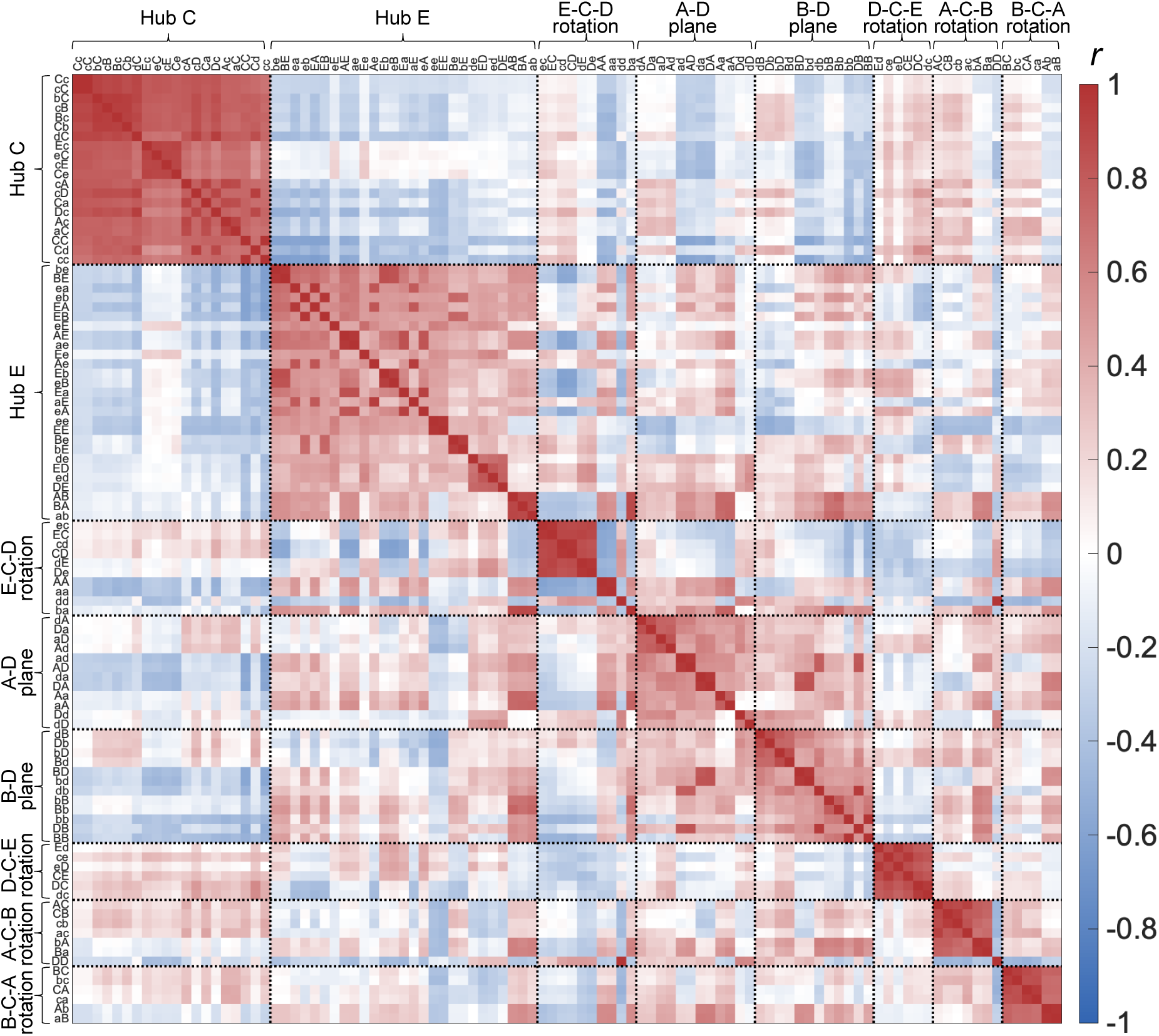
Inter-participant correlation matrix between the number of transitions between EEG microstate templates in Study 1. We divided the EEG topographical dynamics into 100 distinct transitions by applying ten polarity-sensitive EEG microstate templates. Correlations between transition pairs were then calculated based on the number of transitions counted from each participant’s EEG data. The rows and columns were sorted according to the results of Varimax-rotated PCA, in order of absolute loading values, for each of the eight components formed by collecting the items with the highest (most contributing) absolute loading values. Dotted delimiters indicated the boundaries of the components. Each of the eight blocks of diagonal components surrounded by dotted lines corresponded to eight transition components found in the rotated PCA. For simplicity in the axis labels in the figure, positive polarity templates were denoted by uppercase letters (A, B, C, D, E) and negative polarity templates by lowercase letters (a, b, c, d, e), in “transition from” state/“transition to” state order (e.g., “Ab” label meant the transition from A+ to B-).

**Figure 8.**
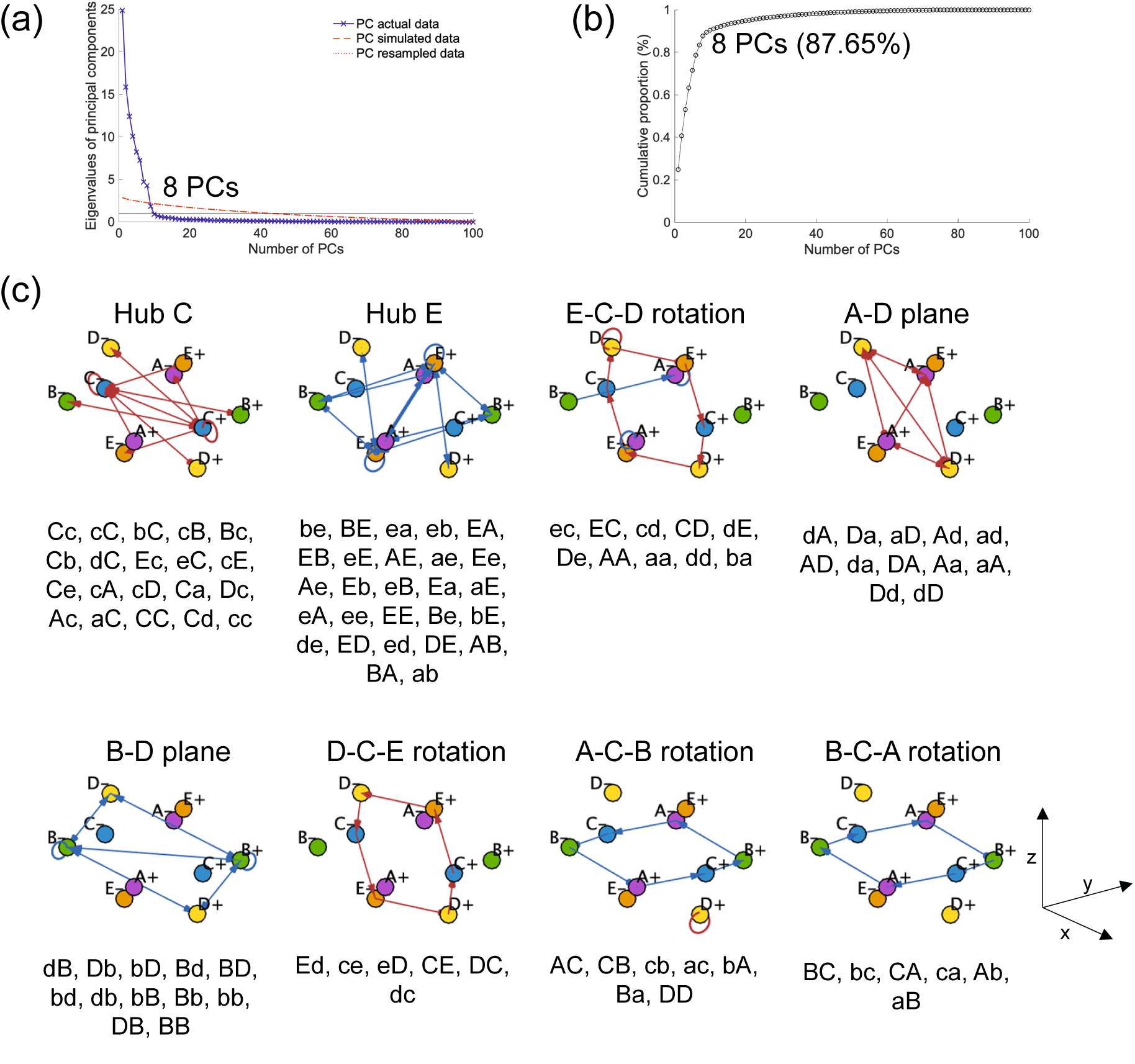
Principal component analysis for EEG microstate transition signatures in Study 1. (a) Parallel analysis scree plot to determine the number of principal components (PCs). The blue line indicated PC eigenvalues obtained from the actual data, and the red dotted lines indicated that they were estimated from simulated/resampled data. Parallel analysis suggested that eight components were the optimal number of PCs. (b) Graph of cumulative contribution ratio as a function of the number of PCs. The eight PCs selected based on the parallel analysis suggestions explained 87.65% of the data variance. (c) List of transition pairs contributing to each PC. After Varimax rotation of the eight principal component axes resulting from the PCA for interpretation, each transition pair was assigned to the component with the highest absolute rotated component loadings. See Supplementary materials (Table S1) for the complete rotated component loading matrix.

Further, we conducted a multivariable logistic regression analysis to examine whether these eight transition component scores predicted binomial age group (0 = younger, 1 = older). For transition components where most transition pairs had negative loading values, the rotated component scores were reversed (hub E, B-D plane, A-C-B rotation, and B-C-A rotation) to align the relationship between age group transition components. The eight transition component scores for the explanatory variables were composite scores based on Varimax rotated PCA, so multicollinearity did not occur (VIFs were 1.02 to 1.23). The overall model demonstrated good fitness and appropriate explanatory variables were included, as evidenced by a non-significant Hosmer-Lemeshow chi-squared goodness-of-fit (GOF) test (X^2^(8) = 11.52, *p* = .17). The Nagelkerke pseudo-R-squared measure showed that all of the predictors together explained 48% of the variance.

The results of the logistic regression analysis are shown in Table 1. All *p* values on logistic analysis were determined using Wald’s test. The model fit results revealed that “hub E” (odds ratio (OR) = 2.98, *p* < .001, 95% CI = [1.88 4.73]), “A-D plane” (OR = 3.01, *p* < .001, 95% CI = [1.87 4.84]), “B-D plane” (OR = 2.29, *p* < .001, 95% CI = [1.44 3.64]), “D-C-E rotation” (OR = 0.45, *p* = .002, 95% CI = [0.27 0.73]), and “A-C-B rotation” (OR = 2.01, *p* = .002, 95% CI = [1.30 3.11]) were significant predictors of age groups. On the other hand, “hub C,” “E-C-D rotation,” and “B-C-A rotation” components were not significant predictors.

**Table 1.** Logistic regression results on predictors of age group by EEG microstate transition components in Study 1.

| *Variable* | *b* | *SE* | *Z (Wald)* | *p* |  | *OR* | *95% CI* |
| --- | --- | --- | --- | --- | --- | --- | --- |
| (Intercept) | -1.11 | 0.22 | -5.13 | .000 |  |  |  |
| Hub C | 0.11 | 0.19 | 0.60 | .550 |  | 1.12 | [0.78 1.61] |
| Hub E | 1.09 | 0.24 | 4.64 | .000 | *** | 2.98 | [1.88 4.73] |
| E-C-D rotation | -0.23 | 0.21 | -1.09 | .275 |  | 0.80 | [0.53 1.20] |
| A-D plane | 1.10 | 0.24 | 4.54 | .000 | *** | 3.01 | [1.87 4.84] |
| B-D plane | 0.83 | 0.24 | 3.51 | .000 | *** | 2.29 | [1.44 3.64] |
| D-C-E rotation | -0.81 | 0.26 | -3.17 | .002 | ** | 0.45 | [0.27 0.73] |
| A-C-B rotation | 0.70 | 0.22 | 3.14 | .002 | ** | 2.01 | [1.30 3.11] |
| B-C-A rotation | -0.27 | 0.23 | -1.19 | .234 |  | 0.76 | [0.49 1.19] |

*Note.* *b*: coefficient for the constant in the null model; *SE*: standard error around *b*; *Z (Wald)*: Wald Chi-square Statistics; *OR*: Odds Ratio; 95% *CI*: 95% confidence interval. \*\*\* $p < .001$ , \*\* $p < .01$ .

This relationship between transition components and age groups was also confirmed by a multivariate *t*-test (Hotelling’s *T*^2^ test) that examined whether there were differences in each transition component score by age group (Figure 9). We found a significant *T*^2^ effect (*T*^2^ = 102.12, *df* = 8, *p* < .001). Post-hoc analysis revealed that the older group showed the larger transition component score of “B-D plane” (*t*(107.11) = 3.57, *p* < .001, adj.*p* = .003, *g* = 0.57), “hub E” (*t*(88.95) = 4.19, *p* < .001, adj.*p* < .001, *g* = 0.73), “A-D plane” (*t*(88.54) = 4.13, *p* < .001, adj.*p* < .001, *g* = 0.72). Also, larger “A-C-B rotation” (*t*(90.96) = 2.38, *p* = .02, adj.*p* = .08, *g* = 0.41) in the older group was marginally significant. On the other hand, less “D-C-E rotation” in older group was marginally significant (*t*(116.31) = -2.59, *p* = .01, adj.*p* = .055, *g* = 0.40).

**Figure 9.**
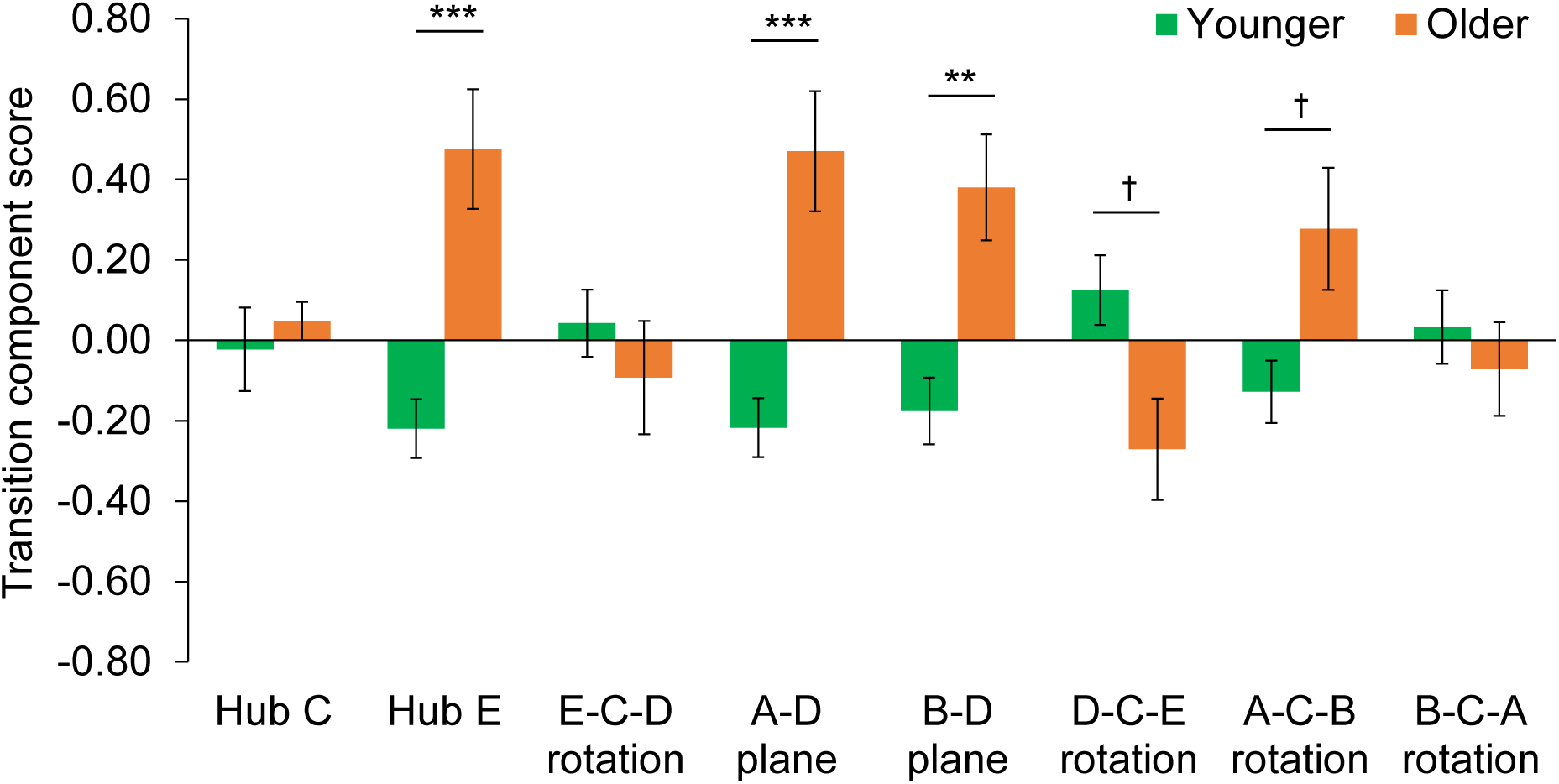
Transition component scores for each age group in Study 1. The green bar shows the transition component scores for the younger group, while the orange bar shows those for the older group. Transition components with mostly negative contributing transition pairs were inverted. Hence, a higher transition component score signifies a pronounced trend in that specific transition component, i.e., the transition probability of the contributing transition pairs tends to be high. Conversely, a lower score indicates the opposite. *Note.* ^***^adjusted *p* (adj.*p*) < .001, ^**^adj.*p* < .01, ^*^adj.*p* < .05, ^†^adj.*p* < .10. Error bars indicated the standard errors.

### 2-3. Discussion

Study 1 updated the EEG microstate analysis by considering topographical polarity. We investigated whether this approach is practical when comparing differences in EEG spatiotemporal transition dynamics between older and younger adults. First, the decreased transitions to C and the increase in A- and B-related transitions in older people shown in previous studies (e.g., Zanesco et al., 2020) were replicated and extended in the present study. Interestingly, these age-related transitions were limited to certain sub-transitions when topographical polarity was considered. Specifically, for the transitions related to microstates C, D, and E, which were less frequent in the older group, a decrease in transitions in the older group mainly occurred between microstates C and D/E of the same topographical polarity as C (especially D± to C± and C± to E±) and between microstates D and E of opposite polarity

(especially E± to D∓). Conversely, for transitions related to microstates A and B, which were dominant in the older group, the increase in transitions mainly occurred within the same topographical polarity and some between polarities. These results suggest that microstate discrete transitions have specific fixed routes and that the frequency of passing among these routes changes with aging.

These transitions within and between polarities depended on the placement of the EEG topography on the neural manifold. If we reformulate the description of EEG dynamics using EEG microstates as a problem of segmentation of the spherical surface and the transition between the segmented areas (Asai et al., 2023), we showed that ten microstates can be placed in a fixed relative position. This finding is not limited to the case of ten microstate maps but can be extended to any number of microstates if the topographical polarity is considered (i.e., the question of how many divisions of the sphere to make). As a result, the distribution of areas corresponding to microstate C was such that other states of the same polarity surrounded them. Such an arrangement limited the transitions involving C in successive two-state transitions to transitions within the same polarity. Also, areas corresponding to A+ and B-, A- and B+, D+ and E-, and D- and E+, where relatively frequent transitions between polarities were observed, were adjacent on the neural manifold space.

The perspective of transitions within and between the topographical polarity, developed by polarity-sensitive fitting, may help describe differences between older and younger groups. The results of this study suggested that between-polarity transitions can occur in only a very limited pattern, although statistical significance could not be determined due to the exploratory nature of the analysis. These limited between-polarity transitions, especially E± to D∓, were less common in the older group than in the younger group. Also, reduced transitions to C have been treated as a characteristic of microstates in older adults, but C-related transitions were limited to within the same polarity. These results suggested that the transition decrease on the axis connecting microstates D-C-E, especially the reduced within polarity D-C and C-E transitions and between-polarity E-D transitions, were characteristic of older people. Furthermore, for A and B-related transitions, the difference between younger and older was in the many transitions among the older between microstates A-D-B-E within the same topographical polarity and between-polarity transition from B to A. Rare transitions were also greater in older adults. Therefore, fewer transitions circling the D-C-E axis, more A-D-B-E within-polarity transitions and B-A between-polarity transitions, and more rare transitions may be biomarkers of aging.

The advantage of considering topographical polarity became more obvious when we took a macroscopic view of transitions from one state to another within or between polarity states. The results of Study 1 indicated differences by age group in the within-polarity transitions from microstates A and C and the between-polarity transitions from microstates B and E. Although our definition of topographical polarity is arbitrary, it appears that both “which country is it currently in (i.e., state)” and “which direction is being travelled (i.e., within and between the polarity)” affect the EEG dynamics with age.

These aging features of microstate transitions were also expressed consistently in the rotated components, separated by transitional features. In this study, discrete transitions were divided into eight components that followed the trajectories on the neural manifold, drawn on the assumption of continuous neural dynamics. These were divided into transitions originating from/to C; transitions originating from/to E; transitions such as moving on a plane containing A and D, as well as B and D; and rotational transitions with a specific direction. Interestingly, among these components, fewer transitions originating from E, fewer transitions between B and D, fewer rotational transitions in the D-C-E and A-C-B directions, and more transitions between A and D were significant predictors of aging.

Among the significant predictors, the component of E-related transitions had the largest explained variance, indicating that transitions connecting to other states with E as a hub are less common in the older group, alongside transitions to C. Microstate E, the fifth state after the four canonical states, may be essential for explaining differences in aging. Although the hub C component did not significantly predict aging, our hub C includes self-recurrences of C and C-related between-polarities transitions, which rarely occur. The fewer transitions to C in the older group are likely reflected in two rotational components, D-C-E and A-C-B rotation. Our findings suggest that the transitions that change with aging are directional. Furthermore, the components capturing transitions between microstate D and passive microstates (i.e., A and B), which are said to be associated with sensory processing (Britz et al., 2010), also changed with aging. These may also reflect a limited hub function of D that was previously unknown. However, these findings are exploratory and need to be validated on another dataset to assess the robustness of these aging features.

## 3. Study 2: Confirmatory analysis of age-related transition differences

Study 2 examined whether the age-related differences in resting EEG state transition dynamics observed in Study 1 represent stable and robust features. To achieve this goal, we examined the reproducibility of the results of Study 1 using independent data.

We applied the updated microstate analysis using our younger-older resting-state spontaneous EEG dataset. Based on the transitions that showed meaningful effect sizes (Hedge’s g) in Study 1, we tested the following hypotheses regarding EEG state transitions in the older group: (a) fewer transitions on the D-C-E axis across polarity, especially fewer within-polarity transitions from D to C and C to E, and between-polarity transitions from E to D; (b) more within-polarity transitions among A, D, B, and E, especially from A to D, D to B, B to A, and between-polarity transitions from B to A; and (c) more rare transitions, especially from E- to D-, D- to A+, E+ to E-, between A±, between B±, and between A-B. When those transitions were crudely categorized into within- and between-polarity transitions, we examined whether they could also be replicated that the older group had more transitions from A± and fewer transitions from C± to within-polarity states than the younger group, as well as more from B± and fewer from E± to between-polarity transitions. We also verified whether the co-occurrence structure of transitions found in Study 1, referred to as transition components, could be confirmed in another dataset. We then confirmed whether those transition components could distinguish age groups, i.e., transitions with E as a hub, transitions connecting passive microstates and D, and unidirectional D-C-E and A-C-B rotation components.

### 3-1. Methods

#### 3-1-1. Participants

Thirty-five healthy adults participated in the experiment. Participants were categorized into two age groups: sixteen younger participants aged 20–40 years comprised the younger group (7 females, 9 males, mean age = 26.0 years, *SD* = 6.0), and 19 older participants aged 61–80 years comprised the older group (9 females, 10 males, mean age = 70.5 years, *SD* = 5.3). All participants gave informed consent before the experiment and provided written consent to participate and share data. This study was approved by the local ethics committee (reference number 19-144 for the Ethics Committee of ATR).

Of the 35 participants, 3 older participants who could not complete all tasks correctly (due to interruption, remeasurement, or repetition of some of the experimental tasks) were excluded from the analysis. Therefore, 32 participants were included in the analysis. The younger group comprised 16 participants (7 females; 9 males; mean age = 26.0 years, *SD* = 6.0; age range, 20–40 years). The older group also comprised 16 participants (6 females; 10 males; mean age = 70.4 years, *SD* = 5.0; age range, 61-79 years).

#### 3-1-2. Experimental settings and procedures

The data were collected as part of another research project on brain function in older individuals, conducted between November 2018 and July 2019. EEG and fMRI data were recorded while participants performed several cognitive tasks on different occasions. During the EEG recording session, participants were asked to perform six different tasks for six minutes per task. The six tasks included four eye-closed tasks (rest, meditation, heartbeat counting, and breath counting) and two eye-open tasks (rest and meditation). The order of the tasks was fixed among the participants. We utilized the eye-closed resting-EEG data in this study.

During the eye-closed resting EEG measurement, participants were instructed to sit in a comfortable chair and remain at rest with their eyes closed, minimizing active thought as much as possible.

#### 3-1-3. EEG data collection, preprocessing, and EEG microstate analysis

EEG signals were recorded using 32 silver-silver chloride electrodes on saltwater sponge-based electrode caps (R-net 32ch, Brain Products GmbH, Gilching, Germany). The 32 electrodes were placed at Fp1, Fp2, Fz, F3, F4, F7, F8, F9, F10, FC1, FC2, FC5, FC6, Cz, C3, C4, T7, T8, CP1, CP2, CP5, CP6, Pz, P3, P4, P7, P8, P9, P10, Oz, O1, and O2, according to the international 10-10 systems. Reference and ground electrodes were placed at FCz and Fpz, respectively. Impedance was kept below 50 kΩ. EEG signals were amplified with an online bandpass of 0.016-250 Hz and digitized using BrainAmp (Brain Products GmbH, Gilching, Germany) at a sampling rate of 500 Hz.

The EEG data underwent the same preprocessing and microstate analysis as those in Study 1. Other than the difference in the data used, no modifications were made to the analysis parameters. However, to avoid the influence of microstate template differences between studies, which is one of the reasons for unstable EEG microstate analysis results (Koenig et al., 2024; Tarailis et al., 2024), we used the same EEG microstate templates in Study 2 created from the LEMON dataset in Study 1. Specifically, we first reduced the template data to 28 electrodes, which were shared electrodes between Studies 1 (61 channels) and 2 (32 channels). The position of each electrode was commonized based on MNI coordinates. Subsequently, the F9, F10, P9, and P10 electrodes, absent in the LEMON data but included in our data, were interpolated using the EEGLAB function “*pop_interp*.” Following this, the EEG microstate templates adjusted for our R-net data were acquired.

#### 3-1-4. Statistical analysis

In Study 2, we confirmed the results obtained in Study 1. The methods for calculating transition probabilities and standardized mean differences were the same as in Study 1. We conducted a multivariate Hotelling’s *T*^2^ test, which tests the difference between two multivariate groups with a chi-square approximation based on each of the following three hypotheses found in Study 1: (a) in older groups, fewer transitions on the D-C-E axis including between-polarity transitions are observed, especially, from D± to C±, C± to E±, and E± to D∓; (b) greater transitions between A-D-B-E (eight within-polarity transitions from A± to D±, D± to B±, B± to E±, E± to A±, and between-polarity transition from B± to A∓); (c) fewer rare transitions from E- to D-, D- to A+, E+ to E-, between A+ and A-, B+ and B-, and A± and B±. Statistical analysis was conducted using RStudio version 2021.09.0 and R version 4.3.1 (R Core Team, 2023). The R functions “*HotellingsT2Test*” and “*effectsize*” were utilized for the multivariate *t*-tests and post hoc Welch’s *t*-test. We applied Holm’s method to adjust the significance level in multiple comparisons. Adj.*p* in the results section represents the adjusted *p*-value, obtained by dividing the conventional significance level of 5% by the number of comparisons for convenience.

To further investigate the robustness of the co-occurrence transition structure found in Study 1, the eigenvectors and the rotation matrix obtained in Study 1 were applied to the data in Study 2. Then, as in Study 1, we reorganized the rotated components based on the maximum loadings of each transition pair. Logistic regression analysis elucidating the age groups and Hotelling’s *T*^2^ test were also conducted using the rotated components obtained in Study 2.

### 3-2. Results

#### 3-2-1. EEG microstate fittings and transitions by considering the topographical polarity

We performed an extended microstate analysis by matching the EEG microstate templates created in Study 1 to the data in Study 2. The results showed that the template maps from Study 1 explained an average of 68.51% (*SD* = 6.29) of the GEV of each participant’s EEG topography time series in Study 2: each microstate label explained 7.42% (*SD* = 2.52) for microstate A+, 5.77% (*SD* = 2.21) for B+, 10.67% (*SD* = 4.98) for C+, 7.54% (*SD* = 4.73) for D+, 3.78% (*SD* = 1.67) for E+, 7.02% (*SD* = 2.37) for A-, 5.56 % (*SD* = 2.06) for B-, or C-, 6.89% (*SD* = 4.07) for D-, and 3.64% (*SD* = 1.51) for E-.

We calculated the transition probabilities between ten states and the differences between each age group. Figure 10 presents the matrices and directed graphs of transition probabilities by older and younger age groups (Figures 10a and 10b), the subtraction results between age groups for each transition pair (older-younger; Figure 10c), and the standardized mean differences between age groups for each transition pair (Figure 10d; also see supplementary Figures S14–16 for additional results using alternative transition matrix definitions).

**Figure 10.**
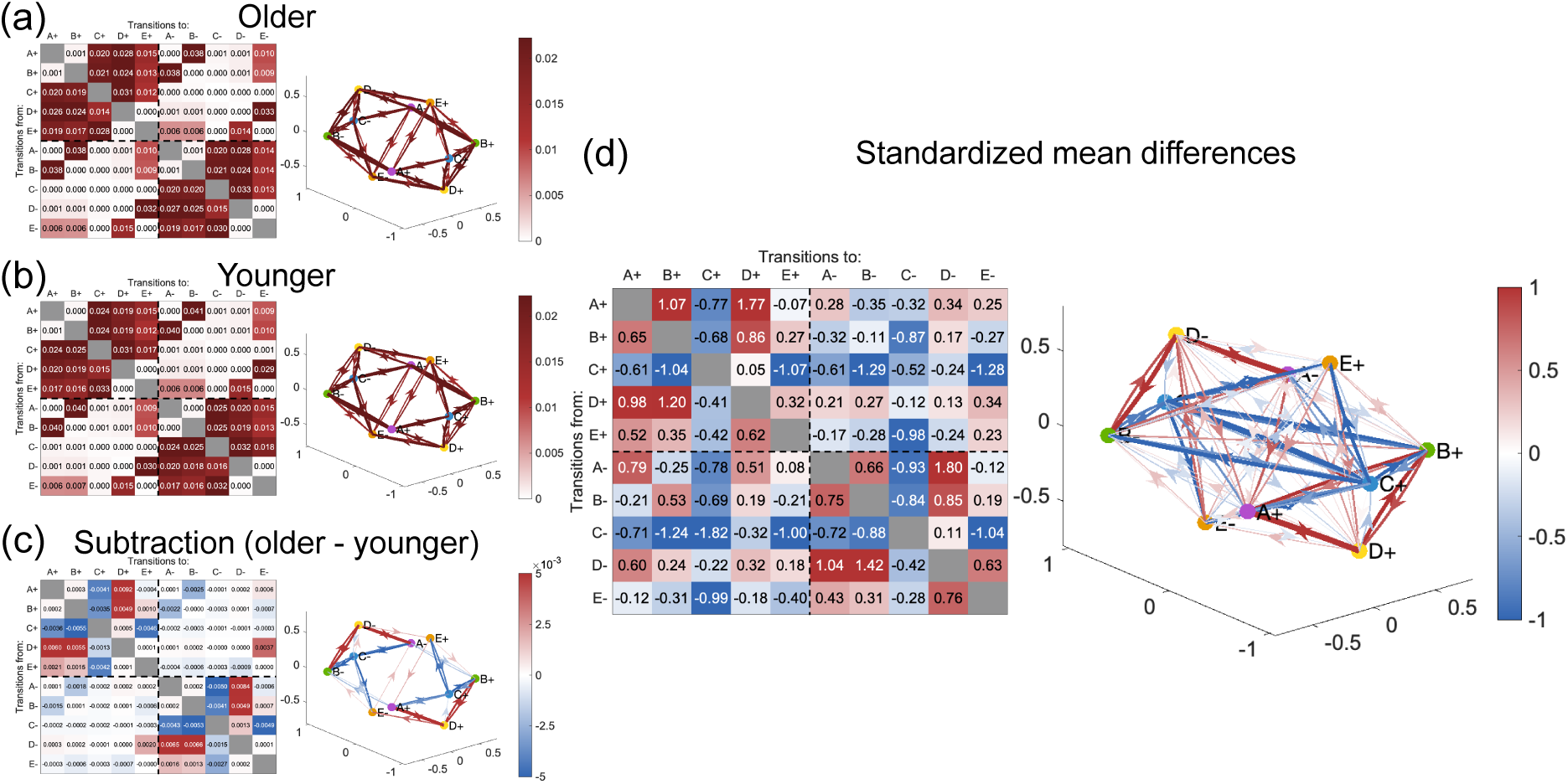
Updated view of microstate transitions—transition probability matrices and directed graphs of microstate transition pairs when considering topographical polarity in Study 2. Each transition pair’s average probability of occurrences was shown by placing the “from” states in the row direction and the “to” states in the column direction. Ninety transition pairs were included, except for ten self-recurrences on the diagonal. In each directed graph, the nodes were the respective template maps, and the edge color indicated the transition probabilities between nodes or the magnitude of the differences between age groups. The edges of the directed graphs were only displayed if the transition probability was greater than 0.011 in (a) and (b) (= the chance level probability defined as random transitions between all states) and greater than 0.0011 in (c) for visualization purposes. (a) Average transition probabilities for the older and (b) younger groups. (c) Differences in the average transition probabilities for each transition pair for older and younger groups. Red indicates more frequent transitions in the older group, and blue indicates less frequent transitions. (d) Standardized mean differences for comparisons between age groups (Hedge’s g; Zanesco et al. (2020)). The symbol of the effect size g and red-blue colors were used to indicate the direction of the difference, with red indicating more frequent transitions and blue indicating less frequent transitions in the older group.

First, we examined whether there was more than one mean difference between the age groups concerning transitions on the D-C-E axis, for which we found promising differences in Study 1. The results showed a significant *T*^2^ effect (*T*^2^ = 16.20, degree of freedom (*df*) = 6, *p* = .01). Post-hoc analysis revealed that some transition probabilities were significantly lower in the older group: the transitions from C+ to E+ (*t*(27.01) = -3.02, *p* = .005, adj.*p* = .03, *g* = 1.04) and C- to E- (*t*(24.97) = -2.95, *p* = .007, adj.*p* = .03, *g* = 1.02).

Second, regarding the hypothesis about transitions among A-D-B-E, we found a significant *T*^2^ effect (*T*^2^ = 52.67, *df* = 10, *p* < .001). Specifically, transition probabilities were more frequent in the older group for the transitions from A+ to D+ (*t*(29.77) = 4.99, *p* < .001, adj.*p* < .001, *g* = 1.72); D+ to B+ (*t*(29.06) = 3.40, *p* = .002, adj.*p* = .01, *g* = 1.17); A- to D-(*t*(28.46) = 5.09, *p* < .001, adj.*p* < .001, *g* = 1.75); and D- to B- (*t*(29.38) = 4.01, *p* < .001, adj.*p* = .003, *g* = 1.38).

Finally, for the third hypothesis about rare transitions, we also found a significant *T*^2^ effect (*T*^2^ = 30.91, *df* = 11, *p* = .001). Post-hoc analysis revealed that the transition from A+ to B+ (*t*(29.63) = 3.02, *p* = .005, adj.*p* = .06, *g* = 1.04) significantly increased in the older group.

In addition, we conducted an exploratory analysis to investigate transitions other than the transition pairs that had promising differences in Study 1. Power analysis using G*Power 3.1.9.7 revealed that an effect size > 1.02 could be detected with an alpha error probability of 0.05, a power of 0.8, and a sample size of 16 participants per group. From Figure 10d, in addition to the hypotheses, several of the transitions from/to C± within and between polarities were less common in the older group: C+ to B+, C+ to B-, C+ to E-, C- to B+, and C- to C+ (|*d*|s = 1.04 to 1.82). Also, concerning the hypothesis about the transitions on the A-D-B-E axis, the transition from D- to A- was more frequent in the older group (|*d*| = 1.04).

In the age-group comparison of the transition probabilities from one state to each of the within- and between-polarity states (Figure 11), based on the results of Study 1, we investigated whether age differences were observed in (d) within-polarity transitions from A±, (e) within-polarity transitions from C±, (f) between-polarity transitions from B±, and (g) between-polarity transitions from E±, using Welch’s *t*-test. For (d), a marginally significant higher within-polarity transition from A+ was observed in the older group (*t*(29.80) = 1.70, *p* = .099, *g* = .59), and no significant difference was found from A- (*t*(30.00) = 1.12, *p* = .27, *g* = .39). For (e), within-polarity transitions from C± were significantly lower for older adults (C+: *t*(28.70) = -3.70, *p* < .001, *g* = 1.27; C-: *t*(27.16) = -3.32, *p* = . 003, *g* = 1.15). On the other hand, for (f) and (g), no significant differences were observed between age groups (B+: *t*(28.35) = -1.11, *p* = .28, *g* = .38; B-: *t*(27.71) = -0.71, *p* = .49, *g* = .24; E+: *t*(29.66) = -0.84, *p* = .41, *g* = .29; E-: *t*(30.00) = -0.76, *p* = .46, *g* = 0.26).

**Figure 11.**
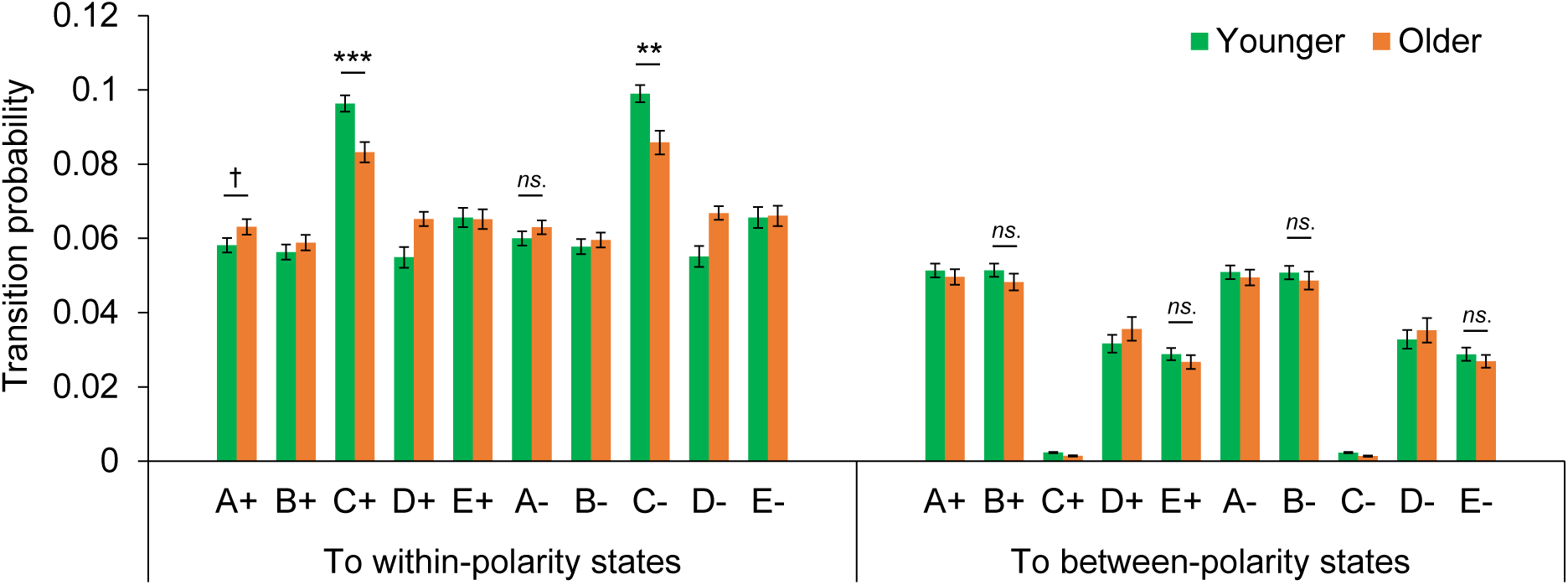
Comparison of transition probabilities from one state to another within and between polarities between age groups in Study 2. The left panel shows the probability of transitioning within polarity from each state (ABCDE±), and the right panel shows the probability of transitioning between polarities from each state. *Note.* ^***^*p* < .001, ^**^*p* < .01, ^*^*p* < .05, ^†^*p* < .10. Error bars indicated the standard errors.

#### 3-2-2. PCA based on the data structure observed in Study 1 and logistic regression analysis for age differences

In Study 2, with 100 transitions, we aimed to assess the contribution of each transition pair to each component in the PC state space identified in Study 1. To achieve this, we first calculated component scores using the eigenvectors obtained in Study 1. Assuming the same eight-dimensional structure as in Study 1, the eight PCs could explain 78.29% of the entire Study 2 data. Subsequently, the rotated component loadings and scores were calculated using the rotation matrix from the Varimax rotation of the eight-dimensional PC space in Study 1 (Table 2). Each transition pair was considered to contribute to the rotated component with the largest absolute loading values. Figure 12 shows the transition components in Study 2 using the weight matrix defined in Study 1. Based on the group of transition pairs contributing to each of the eight rotational components, the eight components were almost congruent with “hub C,” “E-C-D rotation,” “B-D plane,” “hub E,” “D-C-E rotation,” “A-D plane,” “A-C-B rotation,” and “B-C-A rotation” in Study 1.

**Figure 12.**
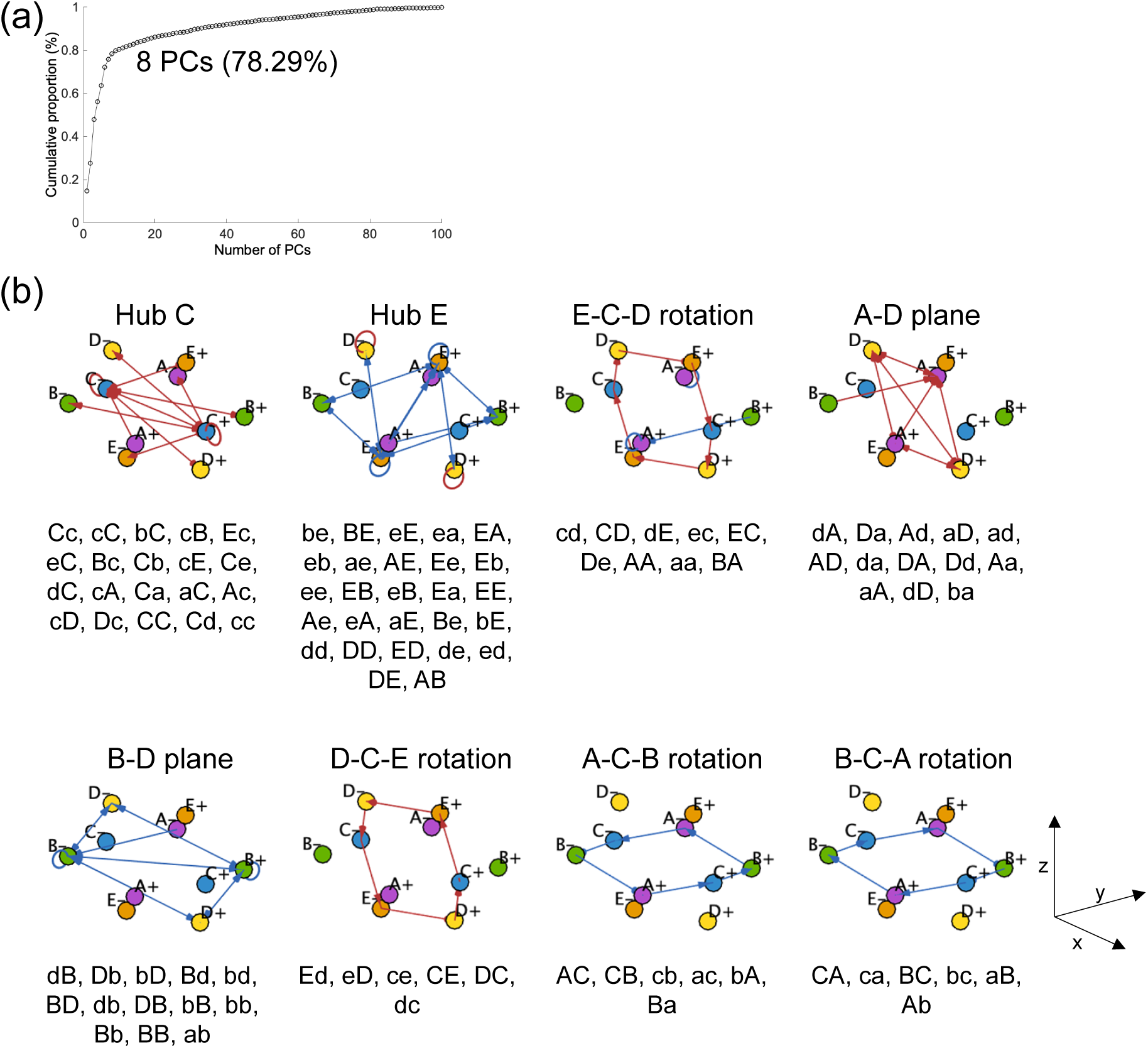
Confirmation of the structure of co-occurrence transitions in Study 2. (a) Graph of cumulative contribution ratio as a function of the number of PCs when applying the eigenvectors and rotation matrix obtained in Study 1 to independent data in Study 2. The eight PCs explained 78.29% of the data variance. (b) List of transition pairs contributing to each PC. After applying the eigenvectors and Varimax rotation matrix in Study 1 to Study 2, each transition pair was assigned to the component with the highest absolute rotated component loadings. Subsequently, we replicated the eight principal component axes. See Supplementary materials (Table S2) for the complete rotated component loading matrix.

**Table 2.**
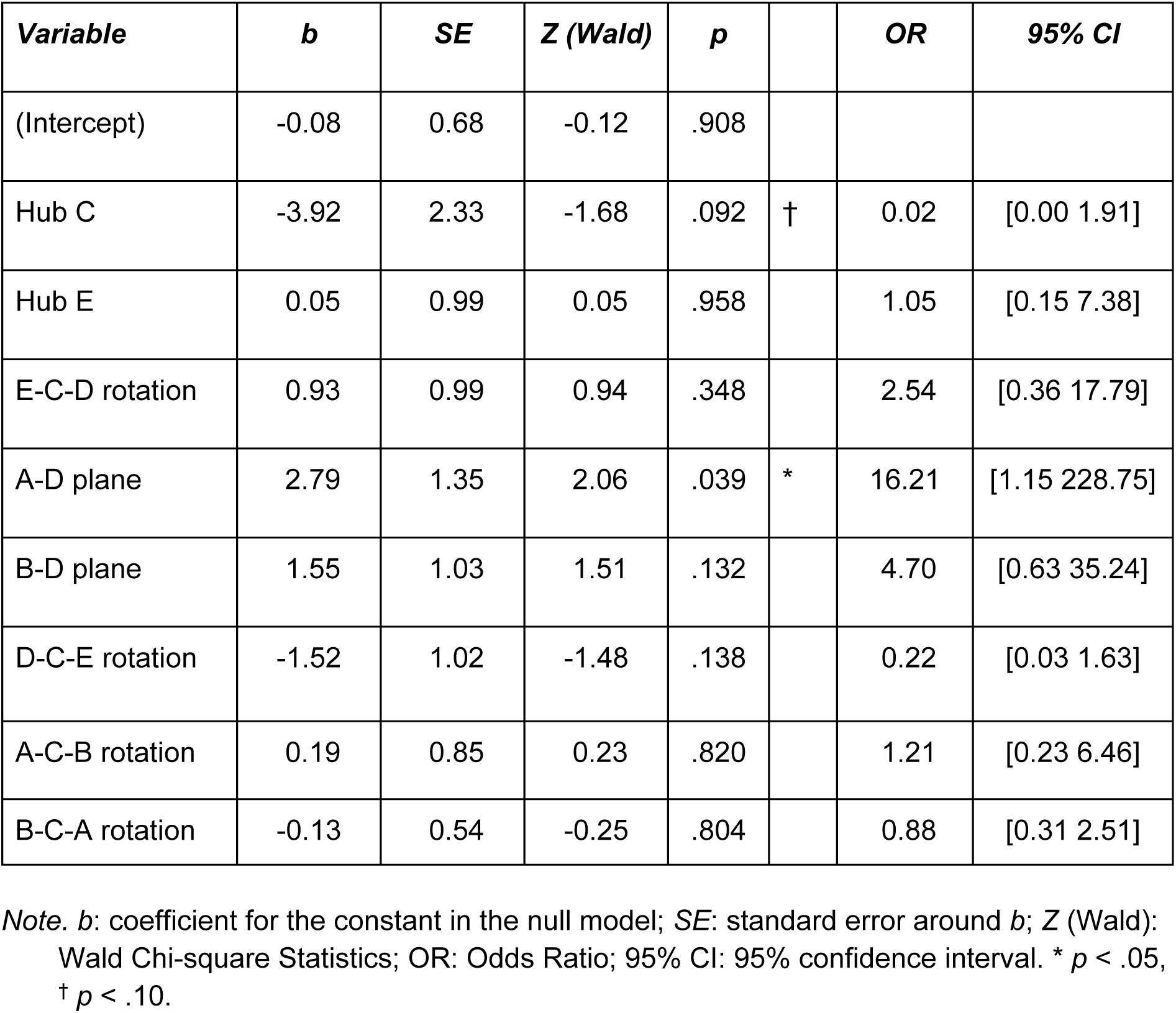
Logistic regression results on predictors of age group by EEG microstate transition components in Study 2.

Furthermore, multivariate logistic regression analysis predicting age groups (0 = young, 1 = old) from the eight transition component scores showed a non-significant Hosmer-Lemeshow chi-square GOF test result (X^2^(8) = 8.60, *p* = .38) for the overall model. This indicates a good fit of the model and the inclusion of appropriate explanatory variables. Nagelkerke pseudo-R-square measures showed that all predictors together explained 73% of the variance. Multicollinearity did not occur because VIF ranged under 10 (1.34–8.74). The logistic regression analysis revealed that “A-D plane” was a significant predictor of age groups (OR = 16.21, *p* = .04, 95% CI = [1.15 228.75]).

The relationship between transition components and age groups was confirmed by Welch’s *t*-test for each of the five transition component scores, the differences of which were found in Study 1 (Figure 13). The results showed a significant age-group difference in “A-D plane” scores, which was higher in the older group (*t*(27.99) = 2.69, *p* = .01, *g* = .93). The “D-C-E rotation” score was marginally significant, lower in the older group than in the younger group (*t*(29.99) = -1.96, *p* = .06, *g* = .67). On the other hand, no significant differences were found for “Hub E” (*t*(27.93) = 0.75, *p* = .46, *g* = .26), “B-D plane” (*t*(29.40) = 0.79, *p* = .43, *g* = .27), and “A-C-B rotation” (*t*(29.35) = -1.21, *p* = .24, *g* = .42). We further examined age group differences for the three non-hypothetical components. We found a significant *T*^2^ effect (*T*^2^ = 10.23, *df* = 3, *p* = .02). Post-hoc tests revealed that the “Hub C” score was lower in the older group (*t*(29.98) = -2.75, *p* = .01, adj.*p* = .03, *g* = .95). “E-C-D rotation” (*t*(25.29) = -0.64, *p* = .53, adj.*p* = .53, *g* = .22) and “B-C-A rotation” (*t*(24.43) = -2.04, *p* = .052, adj.*p* = .104, *g* = .70) were not significantly different.

**Figure 13.**
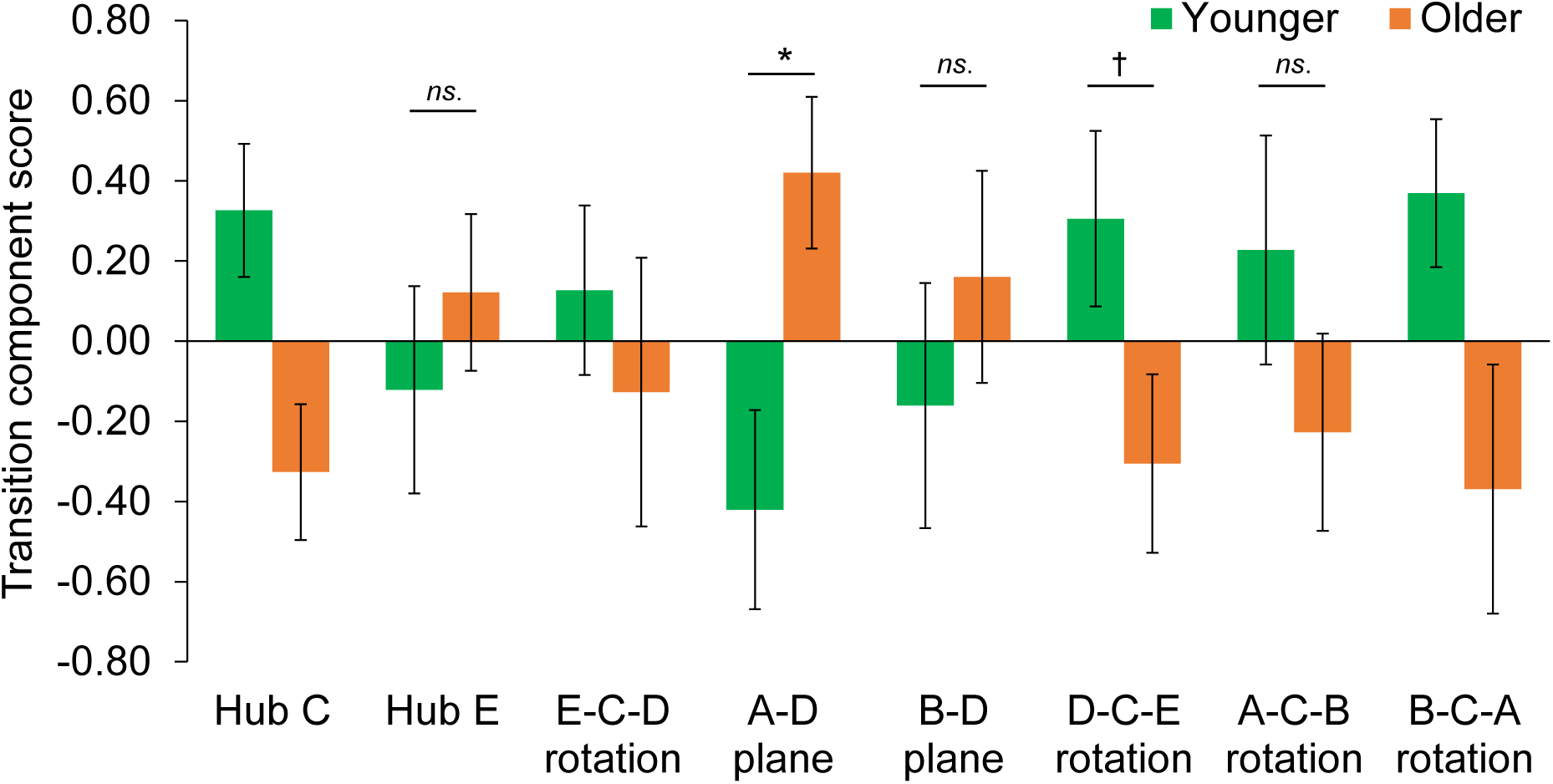
Transition component scores for each age group in Study 2. The green bar shows the transition component scores for the younger group, while the orange bar shows those for the older group. Transition components with mostly negative contributing transition pairs were inverted. Hence, a higher transition component score signifies a pronounced trend in that specific transition component, i.e., the transition probability of the contributing transition pairs tends to be high. Conversely, a lower score indicates the opposite.

### 3-3. Discussion

In Study 2, we attempted to confirm that the EEG microstate features suggested in Study 1 are robust enough to be reproduced using an independent data set. As in Study 1, we performed a polarity-sensitive EEG microstate analysis. We examined three hypotheses on the difference of transition probabilities between age groups based on the results of Study 1: (a) transitions on the D-C-E axis across polarity are less common in the older age group, (b) transitions between A-D-B-E within polarity are more common in the older age group, and (b) rare transitions are more common in the older age group.

The first hypothesis regarding fewer transitions on the D-C-E axis in the older group was partially supported. Based on the observations of Study 1, the transition groups, including transitions from D± to C±, C± to E±, and E± to D∓ were less common in the older group than in the younger group. In particular, within-polarity transitions from D to C and C to E were less frequent in the older group than in the younger group. In contrast, between-polarity transitions between E and D were not significant in Study 2. This suggests that the fixed routes that were unlikely to transition with aging, especially within-polarity transitions between C and D/E at the same polarity, are robust routes that show the effect of aging on transitions.

The second hypothesis regarding the A-D-B-E transition was also supported, as within-polarity transitions from A to D and D to B were more frequent in the older group than in the younger group. The transitions among A-D-B-E were mainly restricted to the polarity, suggesting that the increase in A/B-related transitions found in previous studies may have occurred within this limited polarity. On the contrary, the third hypothesis was not supported, at least in Study 2, and no evidence was presented that the unnatural change in transitions (such as “jumping” between polarities when recorded at a sufficient sampling rate) reflects the effects of aging. The dataset for Study 2 was small but independent of that of Study 1. These results, observed in both Studies 1 and 2, indicated that fewer transitions along the D-C-E axis and more transitions among A-D-B-E within the same polarity are potential biomarkers of aging.

In addition, the results of Study 1 were partially replicated when focusing on transitions from one state to another within the same polarity and between polarities. In particular, the within-polarity transitions from A+ and C± are likely to be biomarkers of aging. On the other hand, no age-related differences were observed in Study 2 for between-polarity transitions from B and E. Despite the potential for undetected differences due to the low power of detection caused by the small sample size in Study 2, at least the within- and between-polarity transition routes from specific states were modulated by aging. Specifically, fewer transitions from A+ and more transitions from C± to within-polarity states are considered to be characteristic of EEG dynamics that change with age.

Furthermore, this study revealed that individual transitions can be grouped into similar transition units based on their co-occurrence relationships, named transition components. The transition components were a robust structure obtained independently of the dataset. They were divided into hubs at C and E, transition groups moving between A/B and D, and unidirectional rotational transition groups with directionality for the D-C-E and A-C-B axes. The cohesion of these transition components has the potential to capture the global EEG transition dynamics rather than problematizing individual transitions.

These transition components may be related to our cognitive functions, mental or physical states, and characteristics. In the present study, we attempted to explain age differences using eight transition component scores. Among the five significant predictors of age identified in Study 1 (“hub E,” “A-D plane,” “B-D plane,” “D-C-E rotation,” and “A-C-B rotation”), only “A-D plane” was significant across datasets. Nonetheless, considering that the number of events per variable generally needs to be greater than 10 (Peduzzi et al., 1996), the present dataset would have required a sample size of at least 10 for each of the eight variables or at least 80 samples. However, there were only 16 participants in each age group. The logistic regression results of Study 2 should be interpreted with caution. The comparison of mean transition component scores of each age group, which partially replicated Study 1, with “A-D plane” scores being higher and “D-C-E rotation” being lower in the older group, had very large standard errors overall, suggesting that there were issues in terms of statistical power. At this time, we reserve the conclusion that the transition components found in this study may reflect our psychological, bodily, or other aspects. This should be further investigated in future studies.

## 4. General Discussion

This study demonstrated that a state-of-the-art microstate analysis considering topographical polarity was appropriate for elucidating differences in EEG transition dynamics with aging. Using a careful two-step approach, we found that when topographical polarity and relative geography among microstates were considered, the older group showed fewer transitions on the D-C-E axis and more frequent transitions among A-D-B-E. Specifically, within-polarity transitions from a particular state were modulated. Moreover, our study indicates that EEG microstate transitions can be decomposed into transition components based on co-occurrence. Age-related changes in microstate transitions may increase or decrease within a larger dynamic context, termed “topographical polarity” in this study. The extended polarity-sensitive microstate analysis we proposed revealed the spatiotemporal continuous transition dynamics, suggesting the significance of polarity-sensitive fitting in microstate analysis.

The age-related differences in polarized microstate transitions in this study may reflect age-related changes in brain structures and networks. Accumulated studies have shown that aging is not only associated with cognitive abilities (e.g., Hedden & Gabrieli, 2004; Mather, 2010; Spreng & Turner, 2019; Tucker-Drob et al., 2019; Verhaeghen & Cerella, 2002) but also with various changes in the brain (e.g., Aron et al., 2022; Cleeland et al., 2019; Li et al., 2001; Oschwald et al., 2019; Samson & Barnes, 2013). For example, aging changes resting-state functional connectivity (Ferreira & Busatto, 2013; Ferreira et al., 2016; Geerligs et al., 2015; Sala-Llonch et al., 2015) and brain structure (Good et al., 2001; Lebel et al., 2010; Oschwald et al., 2019; Resnick et al., 2003). Previous functional neuroimaging studies have proposed age-related changes in gradient patterns, such as antero-posterior and supero-inferior gradients (Boban et al., 2022). For instance, the posterior-anterior shift in aging is a well-known phenomenon, characterized by reduced occipital activity and increased frontal activity (Andrews-Hanna et al., 2007; Davis et al., 2008; McCarthy et al., 2014). This shift is said to be characterized by functional compensation (Davis et al., 2008; Zhang et al., 2017). Additionally, this shift has been observed in structural changes, such as alterations in white matter (Head et al., 2004), and network dynamics (Andrews-Hanna et al., 2007; Zhang et al., 2017). The age-related modulation of microstate transitions on the D-C-E or A-D-B-E axis observed in this study is only a change in the electric field pattern in the topography of scalp EEG. Therefore, it cannot be matched to brain regions observed in imaging studies. However, the well-known structural and network constraints on the direction of information processing in fMRI studies might be manifested as more or less transitions in the microstate transitions on the D-C-E and A-D-B-E axes.

The transitions on the D-C-E axis, which were suggested to be modulated by aging in this study, are state transitions along the midline of the scalp EEG. Studies of age-related EEG changes have often reported findings of age-related attenuation of frontal midline theta activity. Such changes in midline EEG oscillation may be related to aging (Tóth et al., 2014). Our results extend these findings to suggest that EEG oscillation in the midline region may be modulated not only in the time domain but also in the mass domain in relation to aging. Our results pertain specifically to the pattern of supratentorial EEG. Examining the relationship with the A-P gradient, as identified in brain imaging studies, would be valuable in future research. Regarding electrophysiological neural aspects, researchers have primarily concentrated on local changes in specific channels or frequency bands, as well as changes in the time direction. For example, it is known that the alpha oscillation of resting EEG becomes slow, and its power decrease with age (Babiloni et al., 2006; Chiang et al., 2011; Ishii et al., 2017; Klimesch, 1999; Merkin et al., 2023; Scally et al., 2018; Tröndle et al., 2023). Additionally, changes in EEG signal complexity have been observed with aging. Some studies have reported increased complexity in older adults (Anokhin et al., 1996), while others have reported reduced complexity (Lipsitz & Goldberger, 1992; Zappasodi et al., 2015). However, the direction of change remains a topic for discussion due to the diversity of complexity indices (Lau et al., 2022; Ma et al., 2021). Although our results represent only topographical transitions recorded from supracranial EEG, polarity-sensitive microstate analysis allows us to represent spatiotemporal continuous EEG transitions, which may reveal “easy/hard routes to transition” constrained by brain structure and network proximity. Recent discussions in cognitive neuroscience have shifted from the Sherringtonian approach, which elucidates the function of individual brain regions and networks based on functional classification of neuronal populations, to the Hopfieldian approach, which understands the neural basis of cognition from the transition dynamics of masses encompassing local activities (Asai et al., 2022; Barack, 2021; Barack & Krakauer, 2021; Ebitz & Hayden, 2021). The polarized microstate approach may be helpful in such a paradigm shift at the explanatory level.

Our results suggest that the polarity-neglected fitting traditionally employed in microstate analysis may not be appropriate. Traditionally, two topographies of opposite polarity have been considered due to the reversed activity of the same dipole and have been neglected when considering the configuration of the electric field. Indeed, the results of this study showed that the basic properties of the positive and negative polarity were very symmetrical (e.g., the upper and lower triangular matrices of the transition matrix were symmetrical). In this sense, our findings may support the validity of conventional microstate analysis in terms of evaluating the average tendency of a particular electric field configuration to emerge. However, in terms of discretized continuous transitions, this study suggests that even the same D to E transition may have different frequencies of occurrence and even meanings for the D+ to E+ and D+ to E-transitions. Therefore, our findings argued that the two topographies of opposite polarity should be considered separately. At the very least, polarity-sensitive microstate-fitting is helpful because it provides richer, more data-driven information when we follow the neural dynamics, which are essentially continuous transitions.

The polarity of waveforms has originally been important in EEG research, including event-related potential (ERP) studies. Although “polarity” in EEG microstate studies refers to the positivity or negativity of the overall potential pattern at a given instant rather than the polarity of a specific electrode waveform (referred to as “topographical polarity”), it has been indirectly suggested for some time that polarity may have meaningful in microstate studies as well. For example, Tamano et al. (2022) and Asai et al. (2022) suggested the oscillation of EEG microstate dynamics and the existence of both topographical polarities. Moreover, our observations, as well as those of many studies dealing with EEG topography during tasks not restricted to microstates, indicate that EEG microstates during cognitive tasks seem to be biased toward a particular pole, a pattern we call “negative.” Further studies are needed to examine whether each polarity state reflects something functional.

Another good point of the polarity-sensitive microstate fitting is that it allows us to describe microstate dynamics as transitions on a closed loop system (i.e., sphere) in the space of neural manifolds. Neural activity is essentially continuous, and the polarity-sensitive microstate labeling is a good way to address such continuous dynamics as close to its essence as possible, as a transition between divided domains on the surface of a sphere.

This study had several limitations that necessitate further investigation. First, defining topographical polarity in this study posed a challenge. For convenience, the present study referred to the positive microstates as patterns in which the frontal electrode group exhibited positive and the occipital electrode group had negative potentials. Although this was an arbitrary labeling, it is synonymous with calling bipolar global attractors that consequently move between + and - based on the state distribution on the neural state space and transition probability results. The present study suggests an increase in transitions limited within polarity in older adults and limited transition routes across polarity. However, what function this topographical polarity may have and what the transitions between + and – reflect remains unclear. Further accumulation of findings on this point is awaited.

Second is the issue of the robustness of the transition components as reflecting cognitive functions and mental states. In this study, we found eight transition components based on the co-occurrence of 100 microstate transitions. However, it remains unclear how stable these components are regardless of the technique and what the dynamics among transition components are. In this study, the co-occurrence structure of transitions found in a large public dataset acquired in Western countries is at least confirmed when the same component axes were applied to a dataset acquired in Japan. However, it is uncertain whether this structure will consistently emerge in Varimax rotated PCA. Further investigation is needed to determine if these coherent units have any functional significance.

Previous studies on the functional roles of EEG microstates have primarily focused on the functions associated with individual microstates. However, adopting the Hopfieldian perspective, which attempts to explain cognition from the field dynamics, it may be worthwhile to focus on the dynamics of transitions between states and even more abstract transition components rather than the state of EEG microstates to explain cognition.

In summary, the present study revealed age-related differences in EEG transition dynamics, using microstate analysis updated to account for topographical polarity. The transition dynamics associated with aging were characterized by fewer transitions on the D-C-E axis and more between-polarity transitions among A-D-B-E, as well as more transition components associated with the A-D plane. The approach proposed in this study, incorporating topographical polarity, may be useful for separating the conventionally confounded easy and hard transitions and appropriately discretizing the spatiotemporally continuous EEG dynamics.

## Statements and Declarations

### Funding

This work was supported by JST [Moonshot R&D][Grant Number JPMJMS2291], NICT [Research and development of technology for enhancing functional recovery of elderly and disabled people based on non-invasive brain imaging and robotic assistive devices], and the collaboration fund of XNef Inc. and Shionogi & Co., Ltd.

### Conflict of interest

The authors declare that they have no conflict of interest.

### Ethics approval

Study 2 was approved by the Ethical Committee of ATR (approved number: 19-144).

### Availability of data and material

The data in Study 2 are available from the authors upon reasonable request.

## Acknowledgments

We are grateful to Takamasa Hamamoto and Ayako Tsukamoto for their help with data collection in Study 2.

## Authors’ contributions

SK and TA contributed to the study’s conception and design, material preparation, data collection, and analysis. SK wrote the first draft of the manuscript, and TA and HI commented on the manuscript. All authors read and approved the final manuscript.

## Notes

### Competing Interest Statement

The authors have declared no competing interest.

### Summary of Updates

The legends for Figures 11 and 13 have been slightly corrected, and the resolution of all figures has been improved.

