## Supplementary file for "Topographical polarity reveals continuous EEG microstate transitions and electric field direction in healthy aging"

Supplementary materials

**Supplementary A: Toward proper discretization of EEG dynamics as continuous trajectories on a spherical neural manifold**

Asai et al. (2023) visualized continuous EEG topographical dynamics as continuous trajectories on a spherical neural manifold defined based on dissimilarities between topographies. Our polarized EEG microstate transition approach aims to appropriately discretize such spatiotemporally continuous EEG dynamics.

**Figure S1.** *Example of a participant’s eye-closed resting EEG trajectory plotted on a neural manifold based on Asai et al. (2023). This illustration depicts the EEG data of a young participant sampled at 250 Hz for 3 seconds. To visualize the temporal dynamics more clearly, the animation slows down the playback speed to 20 times slower than real-time (i.e., stretched to 60 seconds). The black line represents the trajectory of state transitions over a duration of 100 ms.*

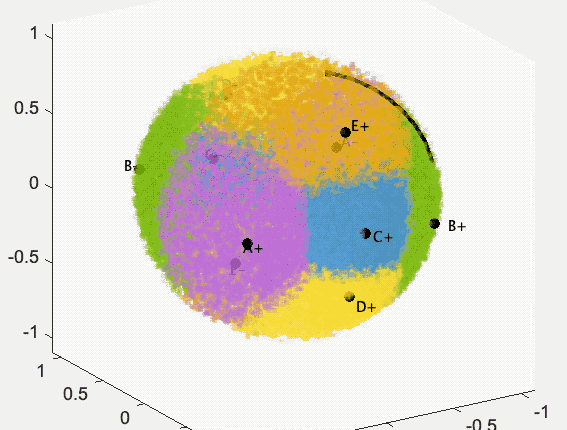

**Supplementary B: Other patterns of transition matrices and directed graphs ignoring topographical polarity in Study 1**

For confirmation analysis disregarding topographical polarity (traditional view), we verified the results of transition matrices and directed graphs for the younger and older age groups using alternative definitions of transition matrices, distinct from those used in the main text. The definitions of transition matrices included the number of transition counts, transition probability with the total number of transitions set to 1, and transition probability to the next state when in a certain state, each with the option to include or eliminate self-recurrence. In the main text, we defined the transition matrix as a transition probability with all transitions except self-recurrences set to 1 (i.e., Figures S3a-c center and right panels and Figure S3d lower panels) because we focused on switching the electric field pattern. Figures S2-4 show when other definitions were employed and when self-recurrence was included or not in each definition.

In each figure, (a) represents the transition matrix and its directed graph for the older group, (b) those for the younger group, (c) the subtraction between the older and younger groups, and (d) the standardized difference between the age groups (Hedge's *g*). In (a)–(c), the left transition matrix shows the case with self-recurrence, the middle transition matrix shows the case without self-recurrence, and the right shows the directed graph without self-recurrence. In (a)-(c) left panels (in the case where self-recurrence was included), since self-recurrences showed tremendous values compared to the others, the color range was determined based on the values of cells other than self-recurrences, and no directed graphs were shown. For (d), the upper and lower panels show the transition matrices and directed graphs when self-recurrence was included and excluded, respectively. In (a) and (b), the edges of the directed graphs were only displayed if the transition probability was greater than the chance level probability. Black squares in each transition matrix indicated transitions with valid edges in the directed graph.

**Figure S2.** *Transition matrices and directed graphs in the case where the transition matrix was defined as the number of transition counts.*

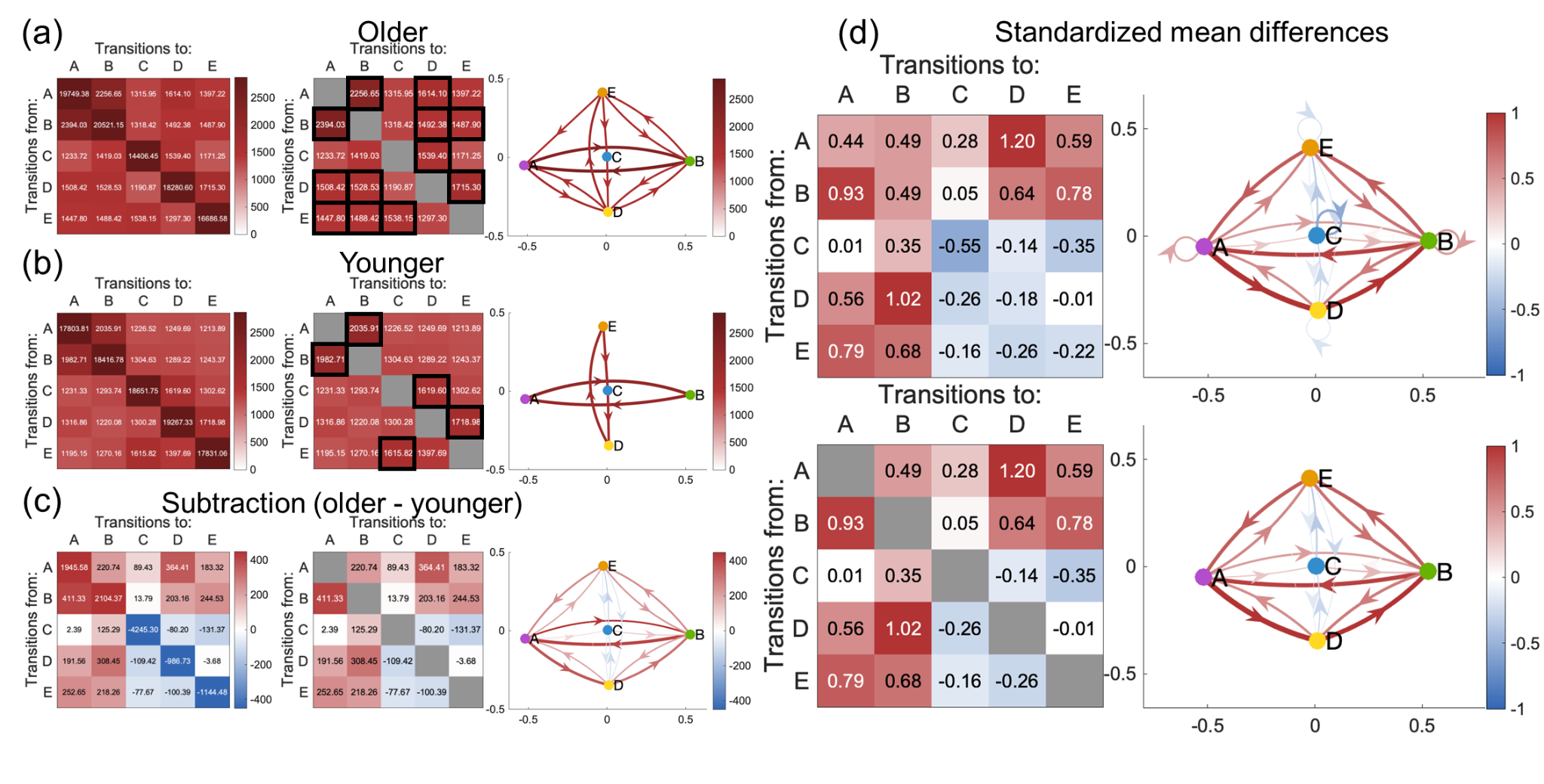

**Figure S3.** *Transition matrices and directed graphs in the case where the transition matrix was defined as transition probability with the total number of transitions set to 1.*

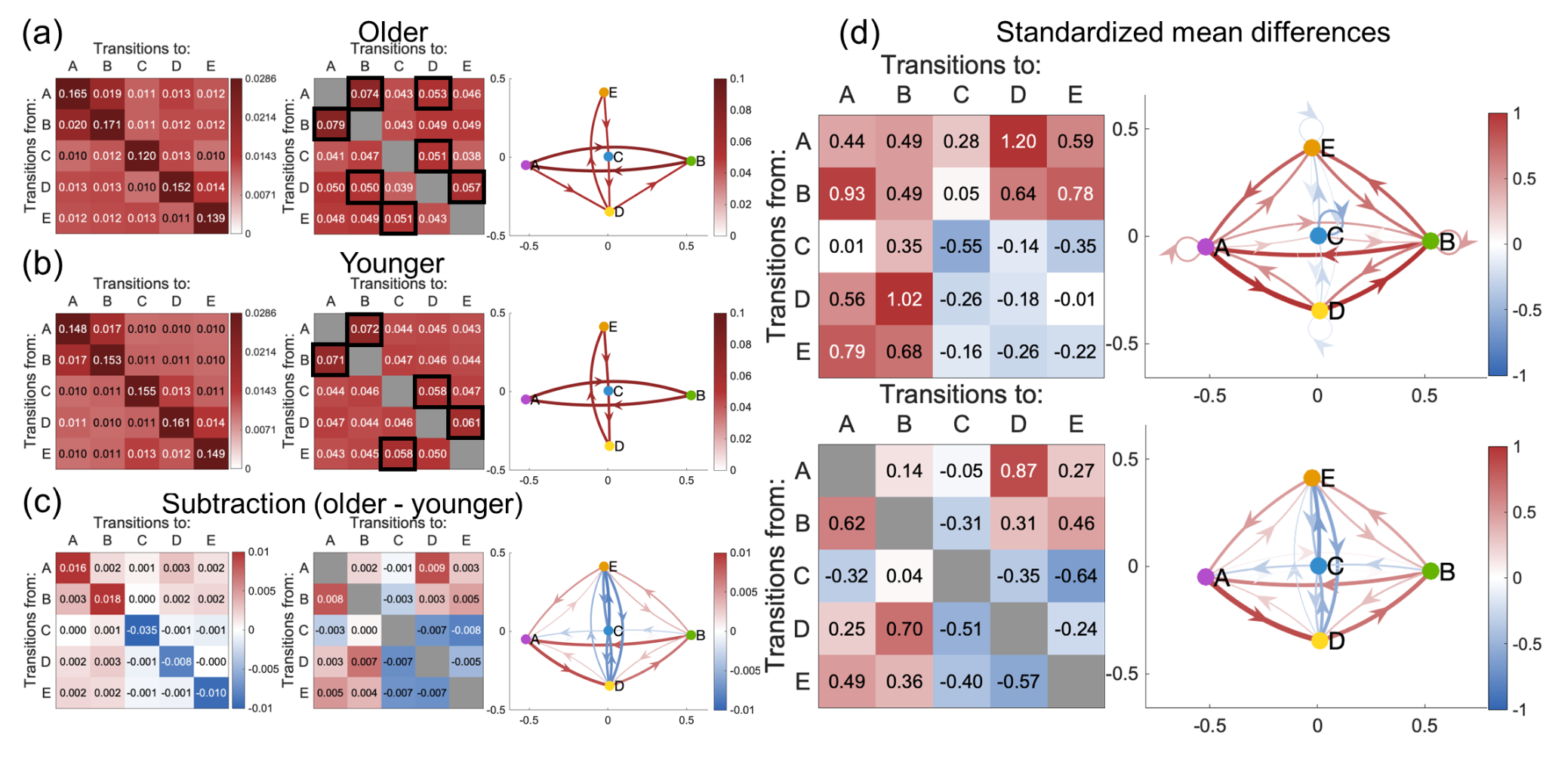

**Figure S4.** *Transition matrices and directed graphs in the case where the transition matrix was defined as transition probability to the next state when in a certain state.*

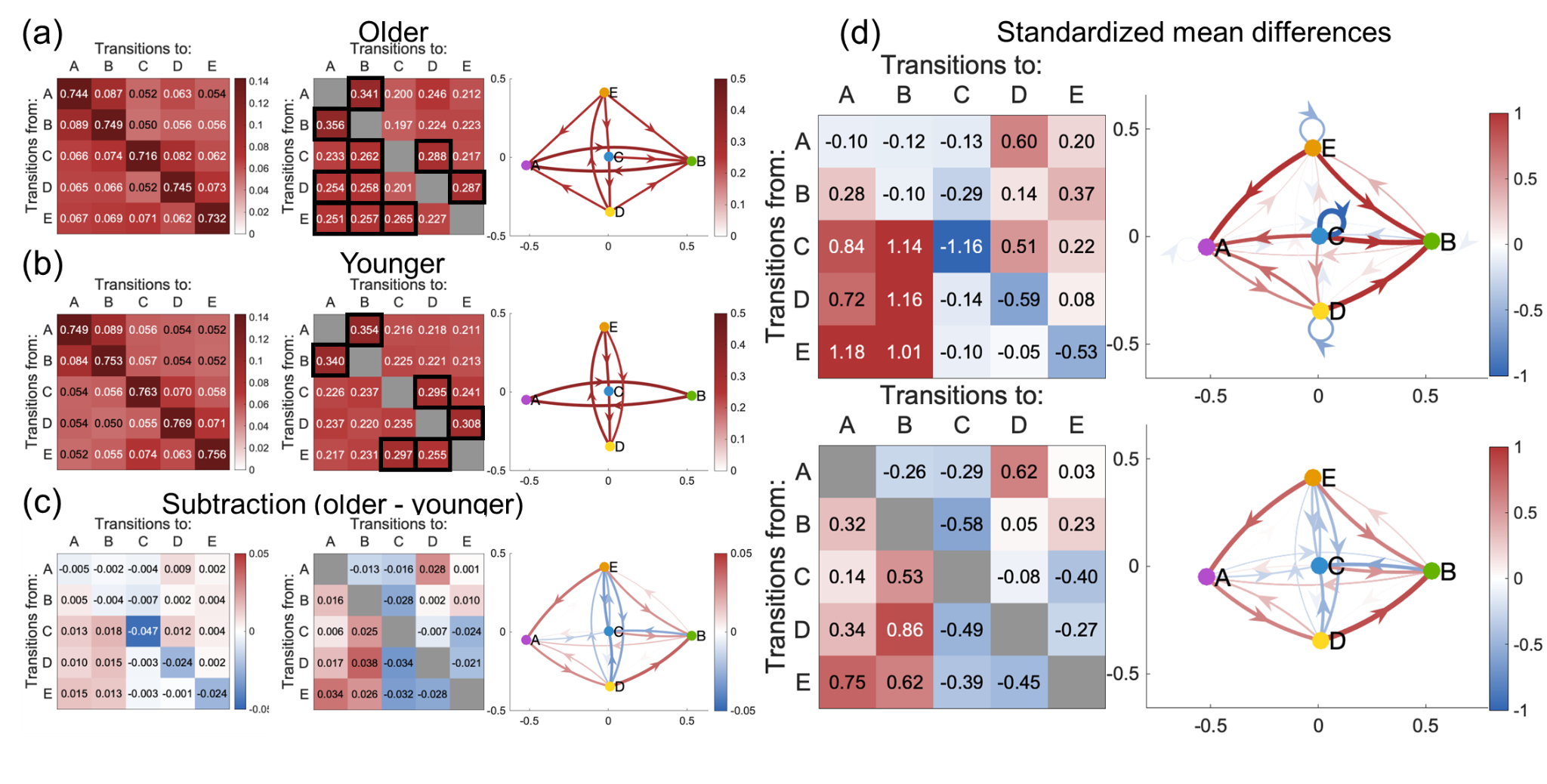

**Supplementary C: 3-D animation of directed graph considering topographical polarity**

When topographical polarity was considered, EEG topographies containing ten microstate templates were aligned on a spherical surface in 3-D state space. Each microstate template map was coarse-grained as a node in a directed graph (see also Figure 1c). Here, we show a 3-D rotation animation of a directed graph to better understand these nodes' relative positions.

It can be seen that microstates ABDE of each pole were arranged in approximately the same 2-D plane, whereas C was not on that plane and was slightly outside of it.

**Figure S5.** *Animation of an example of the 3-D directed graphs of the transitions between 10 microstate template maps considering topographical polarity. Here, as an example, we show a 3-D figure of the standardized difference in transition probabilities between microstates for the younger and older groups of the LEMON dataset (see Figure 4). The distribution of nodes was defined based on the spatial correlation between templates and determined based on the dissimilarity distance compressed by classical MDS (cf., Asai et al., 2023). The color and thickness of the edges in the figure indicate the effect size of family d (Hedge’s g) on the age difference in transition probabilities between its nodes, noting only those edges with a meaningful effect size ≥ 0.44 for visualization purposes.*

**Supplementary D: Other patterns of transition matrices and directed graphs considering topographical polarity in Study 1**

For confirmation analysis considering topographical polarity (updated view), we confirmed the results of transition matrices and directed graphs for the younger and older age groups using alternative definitions of transition matrices, distinct from those used in the main text. The definitions of transition matrices included the number of transition counts, transition probability with the total number of transitions set to 1, and transition probability to the next state when in a certain state, each with the option to include or eliminate self-recurrence. In the main text, we defined the transition matrix as a transition probability with all transitions except self-recurrences set to 1 (i.e., Figures S6a-c center and right panels and Figure S6d lower panels) because we focused on switching the electric field pattern. Figures S5-S7 show when other definitions were employed and when self-recurrence was included or not in each definition.

In each figure, (a) represents the transition matrix and its directed graph for the older group, (b) those for the younger group, (c) the subtraction between the older and younger groups, and (d) the standardized difference between the age groups (Hedge’s *g*). In (a)–(c), the left transition matrix shows the case with self-recurrence, the middle transition matrix shows the case without self-recurrence, and the right shows the directed graph without self-recurrence. In (a)–(c) left panels (in the case where self-recurrence was included), since self-recurrences showed tremendous values compared to the others, the color range was determined based on the values of cells other than self-recurrences, and no directed graphs were shown. For (d), the upper and lower panels show the transition matrices and directed graphs when self-recurrence was included and excluded, respectively.

**Figure S6.** *Transition matrices and directed graphs in the case where the transition matrix was defined as the number of transition counts.*

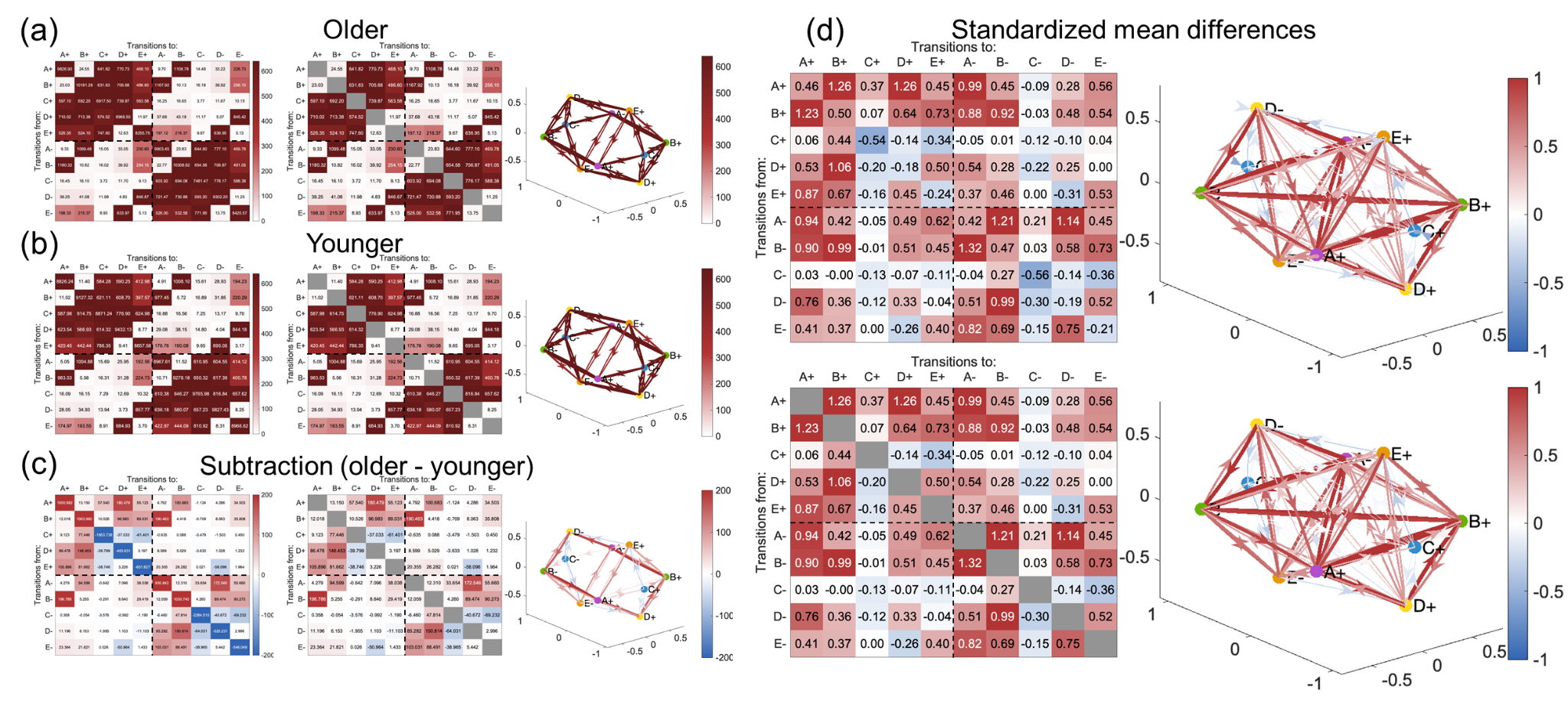

**Figure S7.** *Transition matrices and directed graphs in the case where the transition matrix was defined as transition probability with the total number of transitions set to 1.*

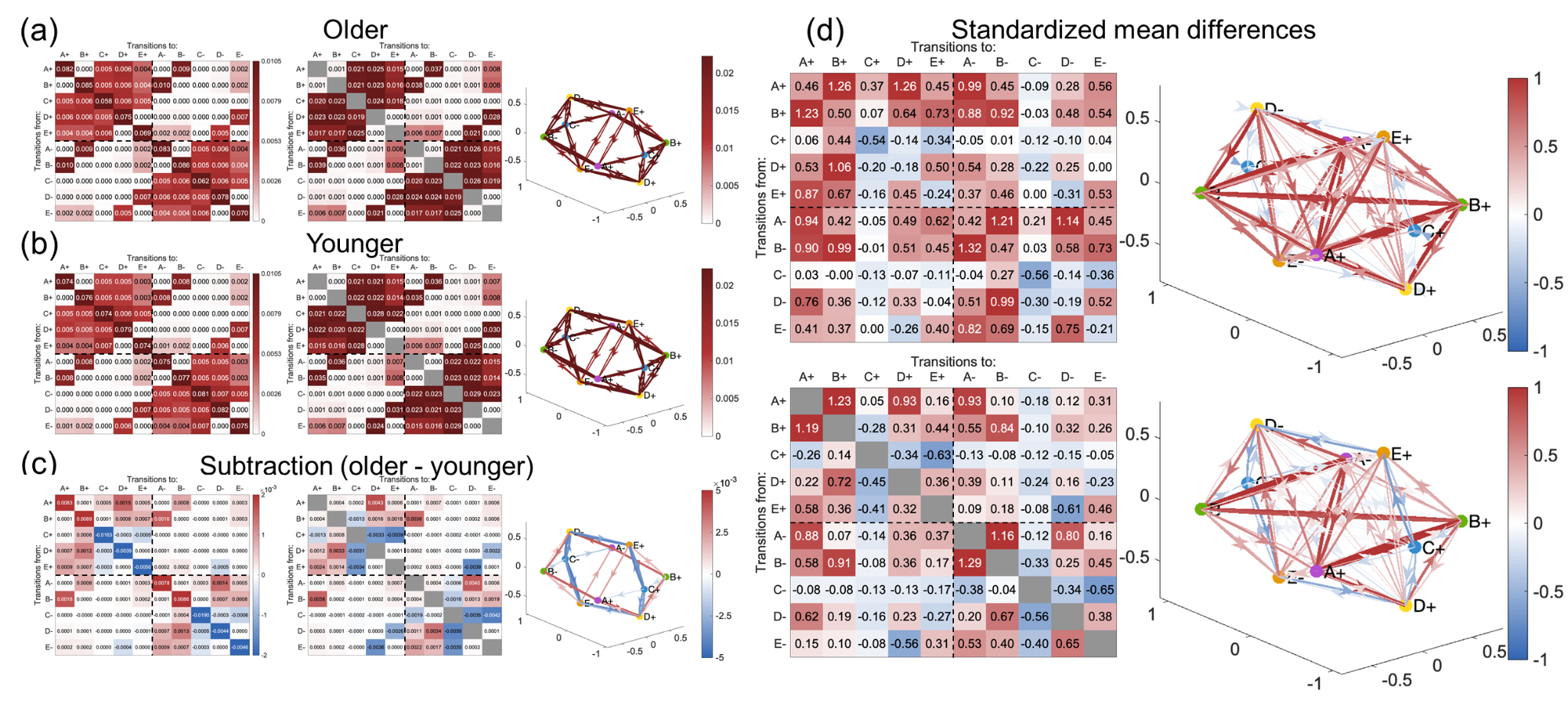

**Figure S8.** *Transition matrices and directed graphs in the case where the transition matrix was defined as transition probability to the next state when in a certain state.*

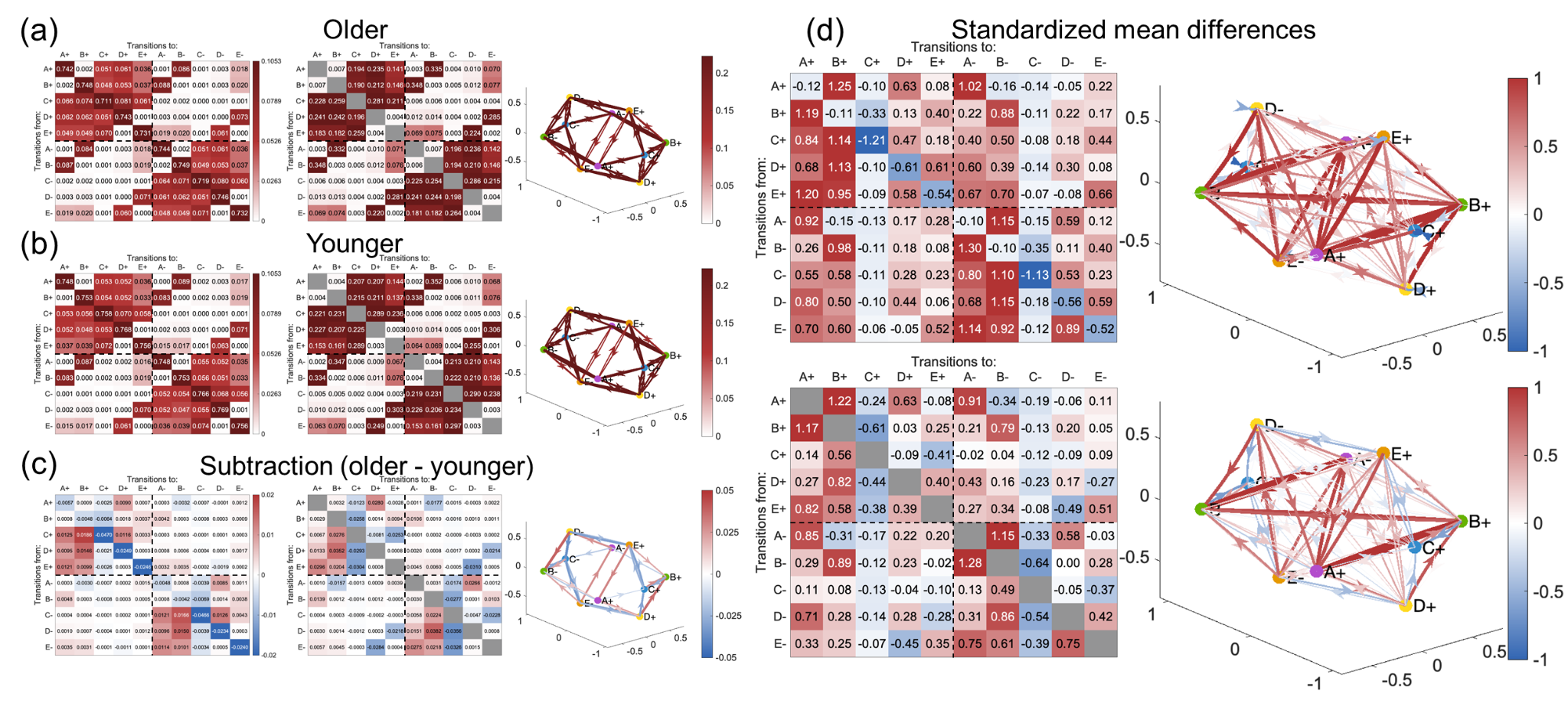

**Supplementary E: Results of Varimax rotated PCA in Study 1**

We conducted a Varimax rotated PCA, employing orthogonally rotated 8 PCs obtained from a PCA of 100 transition pairs. The rotated component loadings for each transition pair are shown in Table S1. Transition pairs were grouped as belonging to the component with the highest absolute value of rotated component loadings (highlighted in yellow in the table). Each transition component was named based on the content of the transition pairs to which it belonged. For convenience, each transition pair was labeled with positive states in upper case (e.g., A+ was denoted by “A”) and negative states in lower case (e.g., A- was denoted by “a”), in order of “from” and “to” states (e.g., the transition from A+ to A- was denoted by “Aa”).

**Table S1.** *Rotated component loadings of each transition pair in Study 1. Loading values were calculated using Varimax-rotated PCA.*

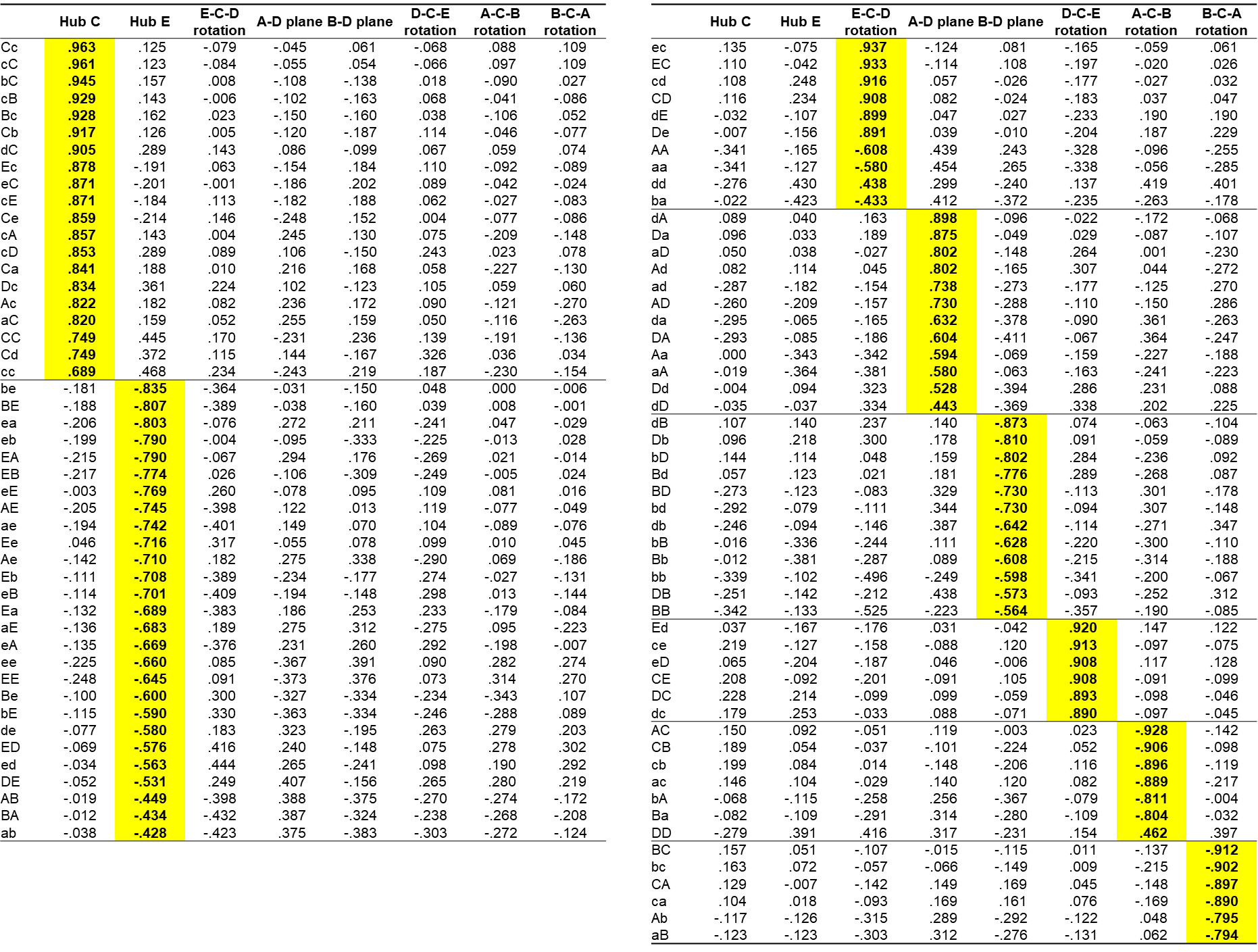

To determine whether transition components clustered based on the magnitude of loadings obtained by Varimax-rotated PCA could be detected stably by other methods, *k*-means clustering and hierarchical clustering (Ward’s method) were also performed. In this process, the transition counts of the participants’ 100 transition pairs were standardized per transition pair.

The results showed that regardless of methods, Hub C and the four rotation components were robustly obtained, confirming a stable transition component structure (Figures S8 and S9). The hierarchical clustering results also showed that the two pairs of transition components (clusters) of E-C-D and D-C-E rotations, and A-C-B and B-C-A rotations, which have different directions on the same axis, were relatively distant from each other and clearly separated by the transition direction even if the transitions take the same route. On the other hand, Hub E and the A-D/B-D plane components were somewhat unstable between cluster methods.

**Figure S9. Eight clusters of EEG microstate transition signatures obtained by the *k*-means clustering.**

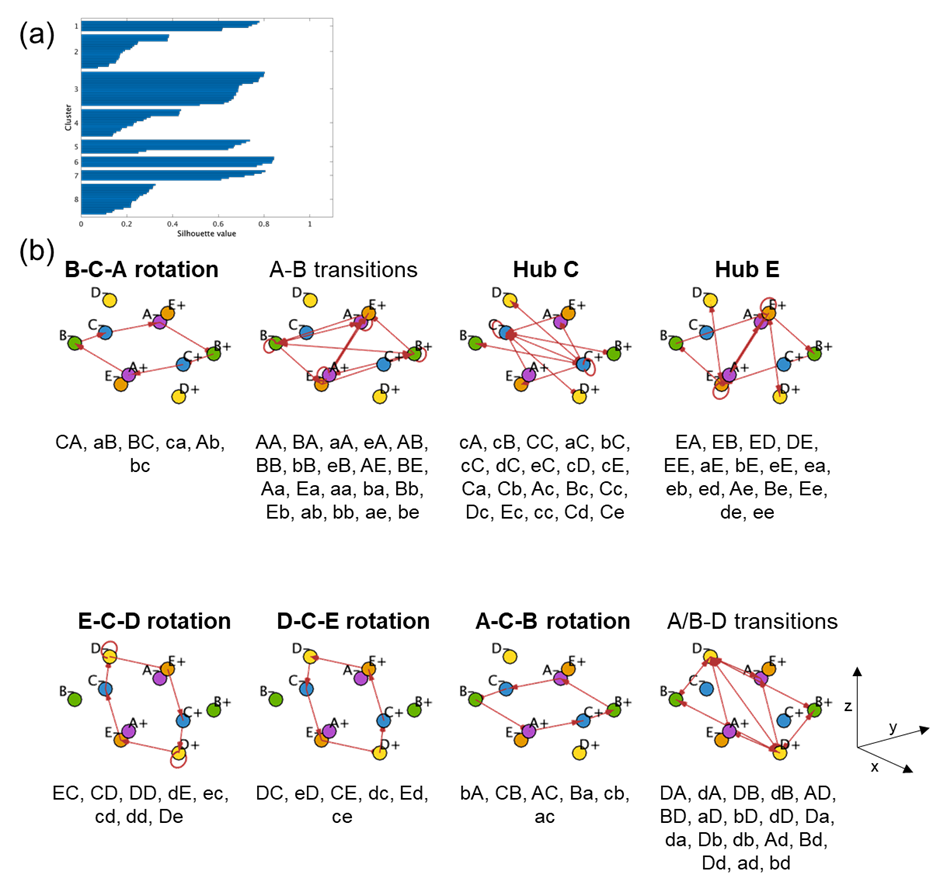
*(a) Plot of silhouette coefficients. Transition pairs comprising clusters 1, 3, 5, 6, and 7 had sufficient silhouette coefficients. (b) Eight transition components discriminated through k-means clustering. We performed k-means clustering with k = 8 under based on 1000 maximum repetitions and 50 iterations. Cluster names in bold indicate clusters in which the most transition pairs belonging to the cluster were the same as those obtained by PCA-based clustering.*

**Figure S10. Eight clusters of EEG microstate transition signatures obtained using hierarchical clustering.**

*(a) Dendrogram of 100 transition pairs. Distances between clusters were calculated using Ward’s method. Label colors indicated clusters separated by more than 40% of the maximum inter-cluster distance. (b) Eight transition components discriminated by hierarchical clustering. The colors of the cluster names corresponded to the colors in Figure S9a. Cluster names in bold indicate clusters in which the most transition pairs belonging to the cluster were the same as those obtained using PCA-based clustering.*

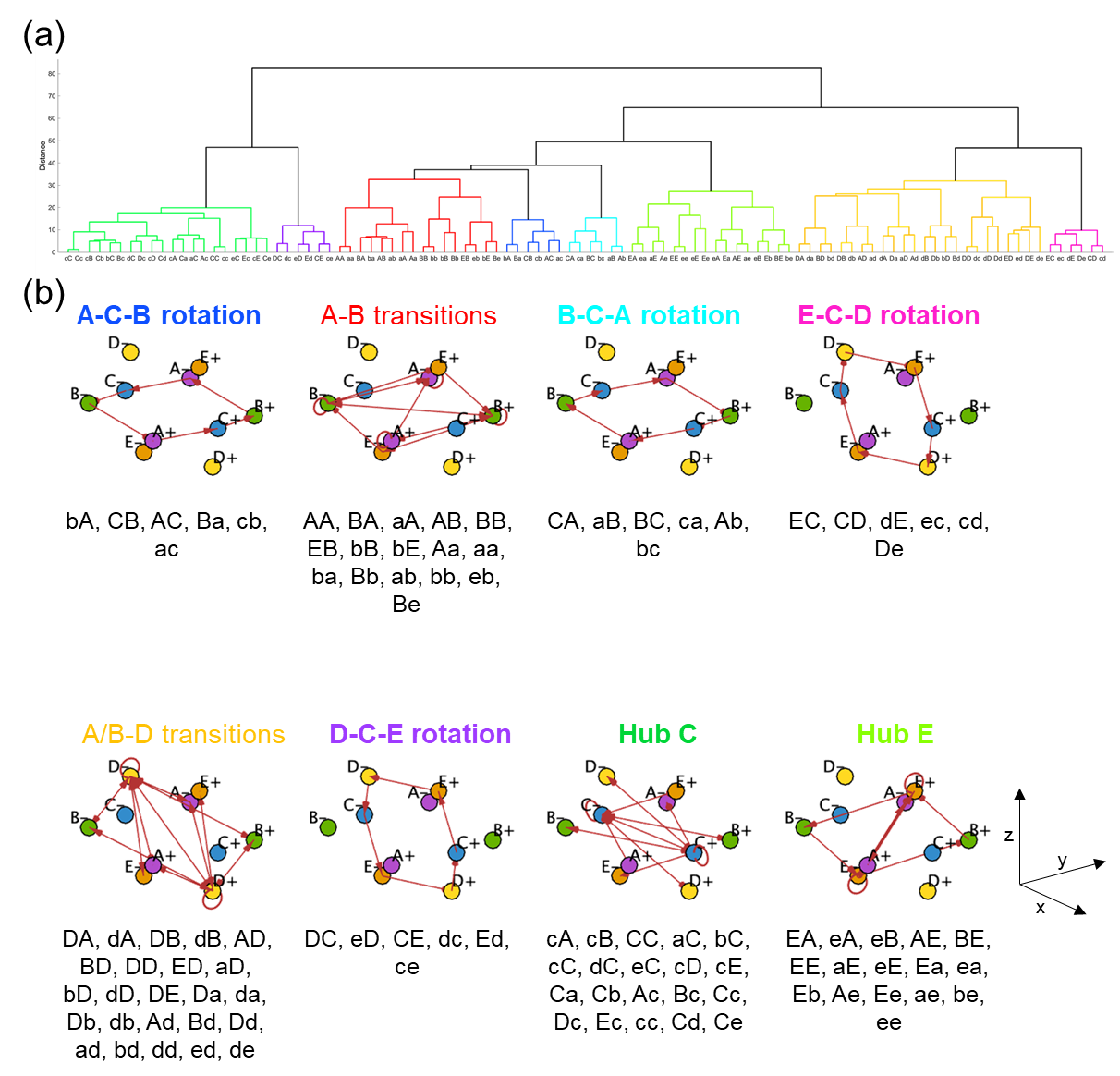

**Supplementary F: Other patterns of results of Varimax rotated PCA in Study 1 when using other definitions of the transition matrix**

We conducted multivariable logistic regression analysis to examine whether the eight transition component scores predicted binomial age group (0 = younger, 1 = older) using other definitions of the microstate transition matrix.

Once we define a transition matrix in a certain way, we can find a cohort of co-varying transitions according to that definition. In this section, we used six different definitions, including three different ways of representing transitions and two that included or excluded self-recurrence, to conduct the rotated PCA and logistic regression analysis.

**Figure S11.** *Varimax rotated component analysis and logistic regression analysis of predictors of age group by EEG microstate transition components in the case where the transition matrix was defined as the number of counted transitions.*

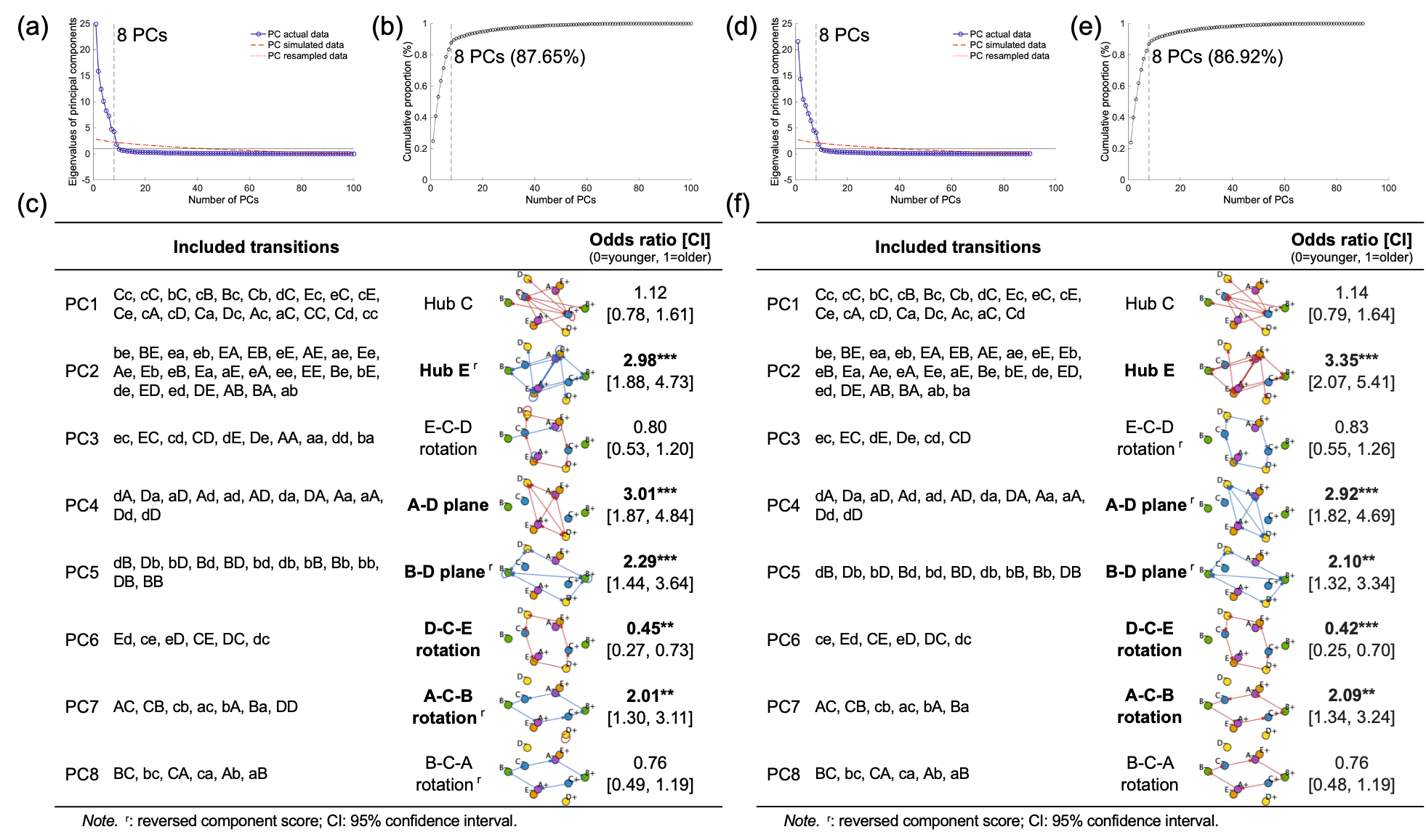

**Figure S12.** *Varimax rotated component analysis and logistic regression analysis of predictors of age group by EEG microstate transition components in the case where the transition matrix was defined as transition probability with the total number of transitions set to 1.*

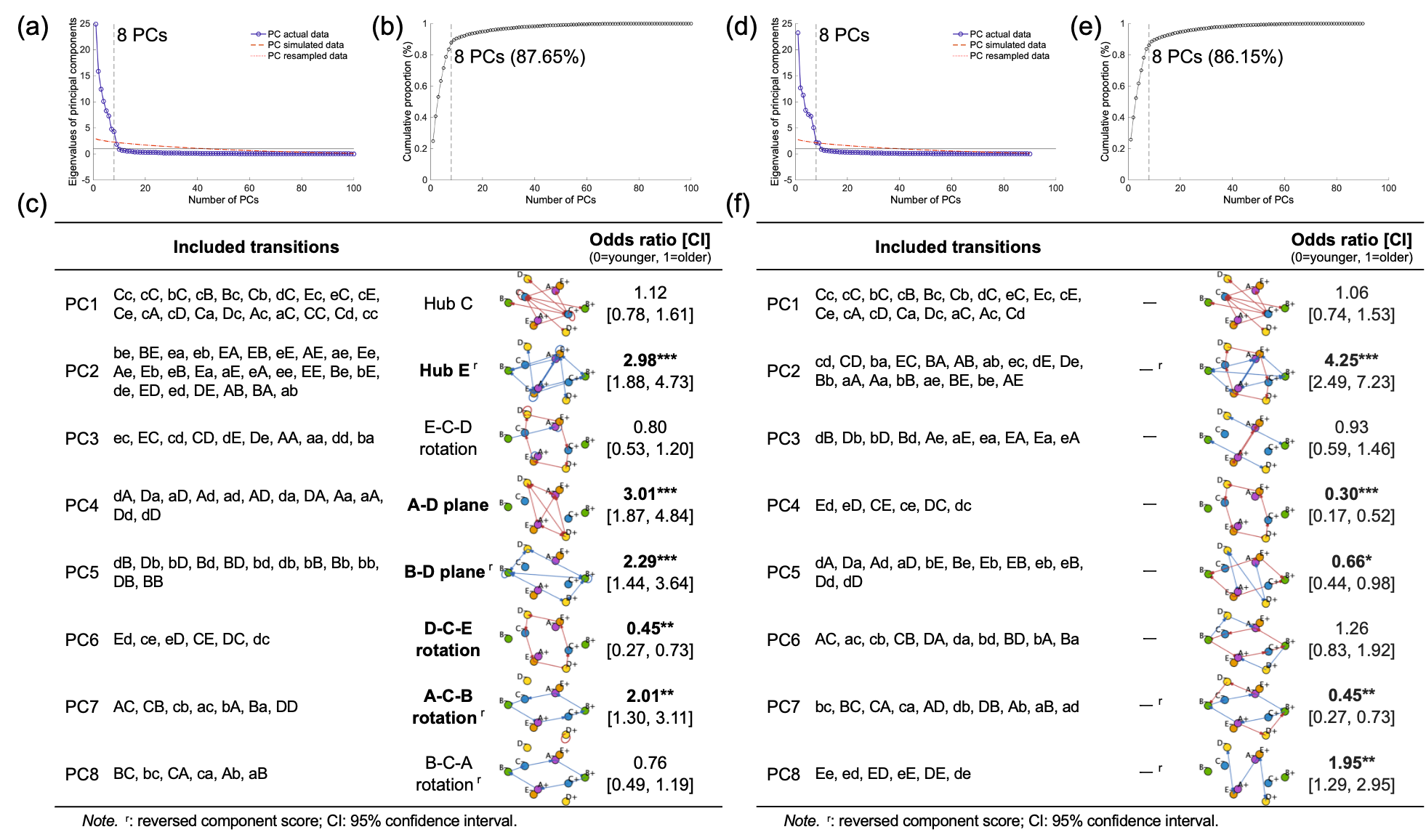

**Figure S13.** *Varimax rotated component analysis and logistic regression analysis of predictors of age group by EEG microstate transition components in the case where the transition matrix was defined as transition probability to the next state when in a certain state.*

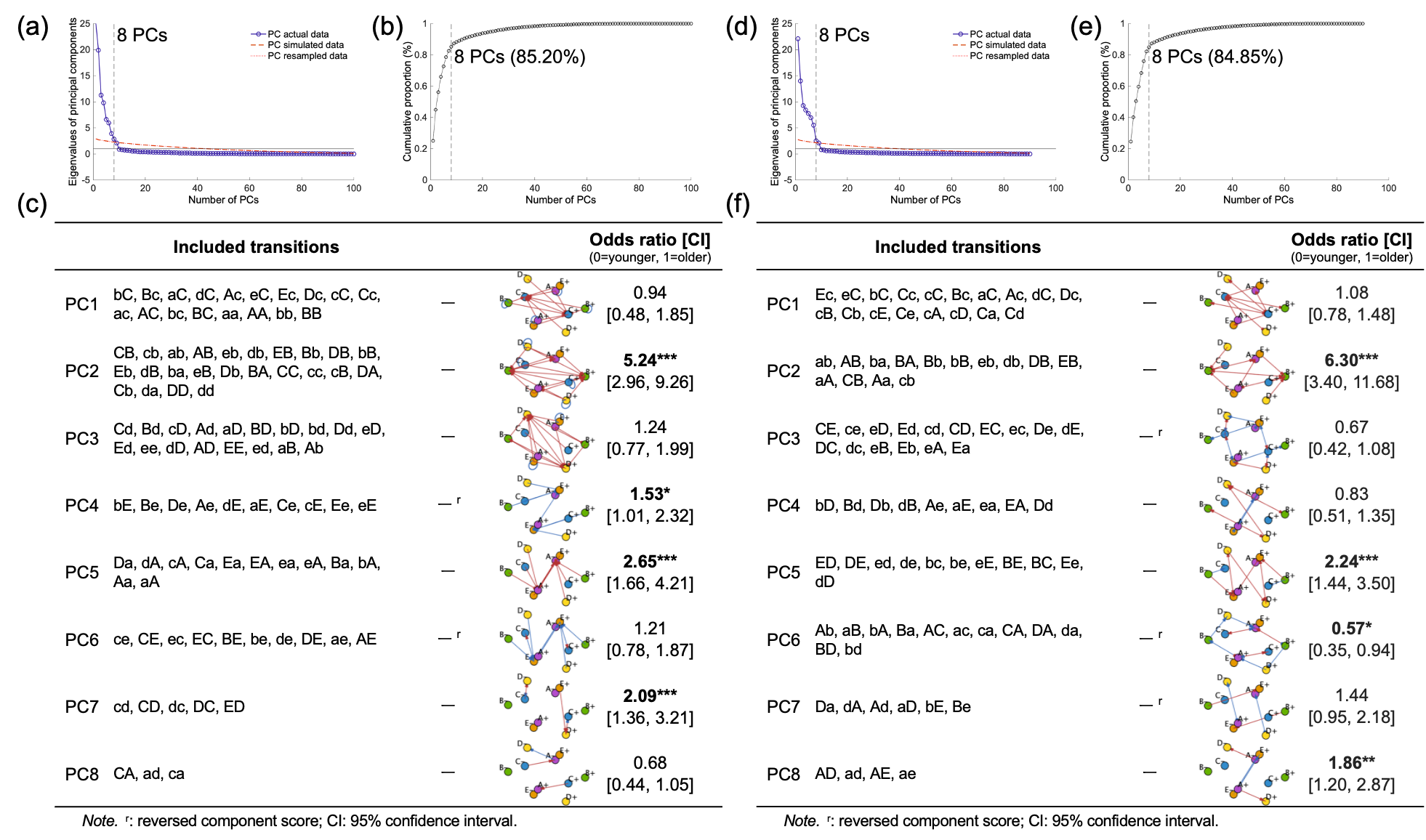

**Supplementary G: Other patterns of transition matrices and directed graphs considering topographical polarity in Study 2**

For confirmation analysis considering topographical polarity (updated view), we confirmed the results of transition matrices and directed graphs for the younger and older age groups using alternative definitions of transition matrices, distinct from those used in the main text. The definitions of transition matrices included the number of transition counts, transition probability with the total number of transitions set to 1, and transition probability to the next state when in a certain state, each with the option to include or eliminate self-recurrence. In the main text, we defined the transition matrix as a transition probability with all transitions except self-recurrences set to 1 (i.e., Figures S14a-c center and right panels and Figure S14d lower panels) because we focused on switching the electric field pattern. Figures S13-S15 show when other definitions were employed and when self-recurrence was included or not in each definition.

In each figure, (a) represents the transition matrix and its directed graph for the older group, (b) those for the younger group, (c) the subtraction between the older and younger groups, and (d) the standardized difference between the age groups (Hedge’s *g*). In (a)–(c), the left transition matrix shows the case with self-recurrence, the middle transition matrix shows the case without self-recurrence, and the right shows the directed graph without self-recurrence. In (a)–(c) left panels (in the case where self-recurrence was included), since self-recurrences showed tremendous values compared to the others, the color range was determined based on the values of cells other than self-recurrences, and no directed graphs were shown. For (d), the upper and lower panels show the transition matrices and directed graphs when self-recurrence was included and excluded, respectively.

**Figure S14.** *Transition matrices and directed graphs in the case where the transition matrix was defined as the number of transition counts.*

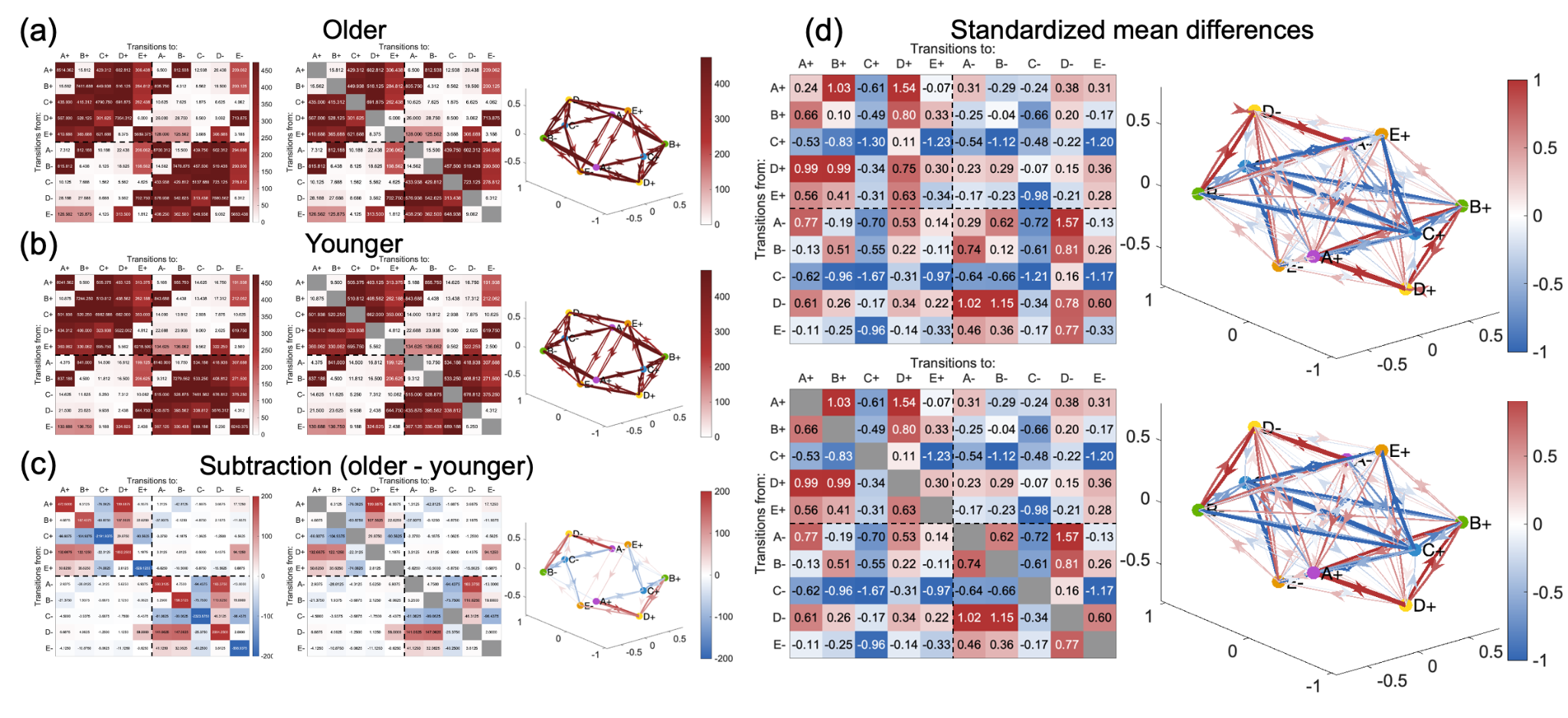

**Figure S15.** *Transition matrices and directed graphs in the case where the transition matrix was defined as transition probability with the total number of transitions set to 1.*

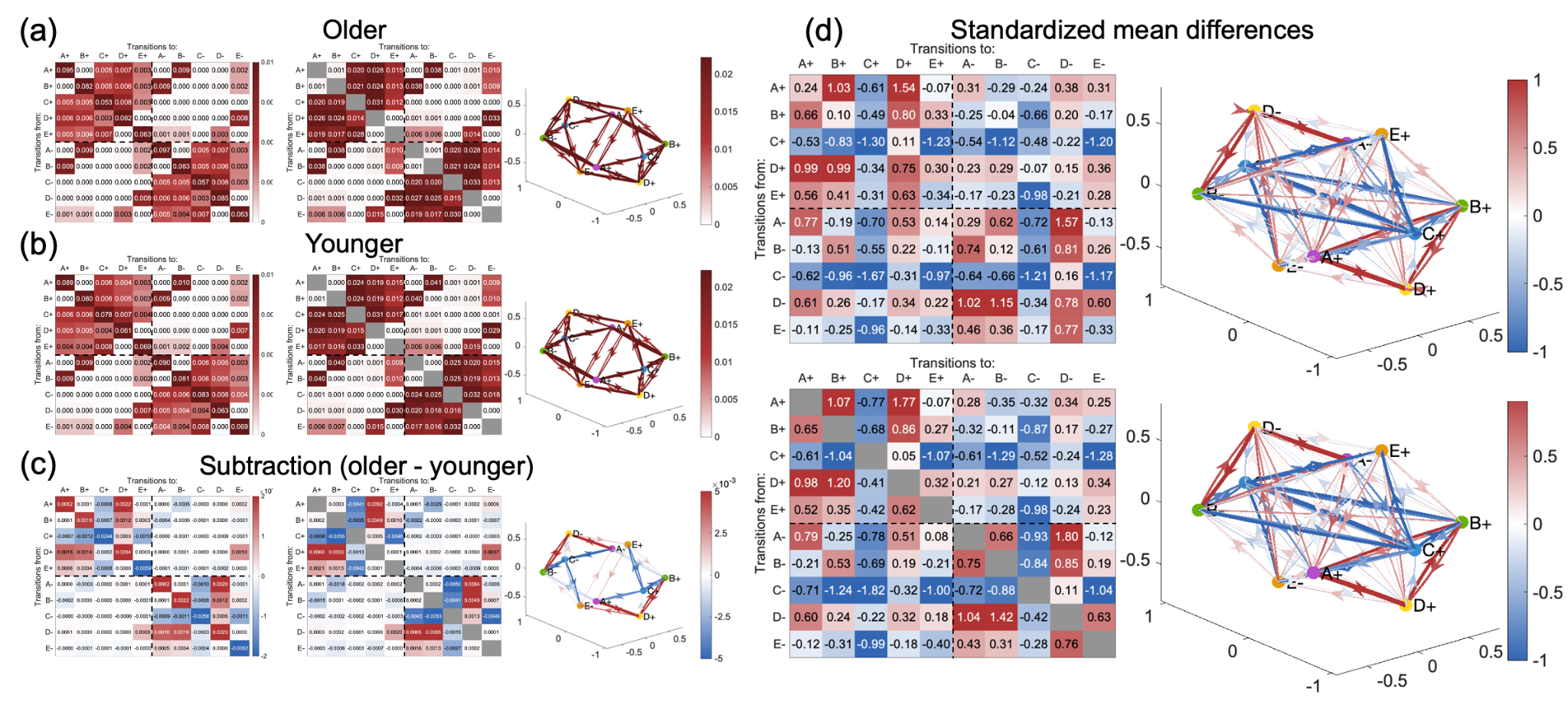

**Figure S16.** *Transition matrices and directed graphs in the case where the transition matrix was defined as transition probability to the next state when in a certain state.*

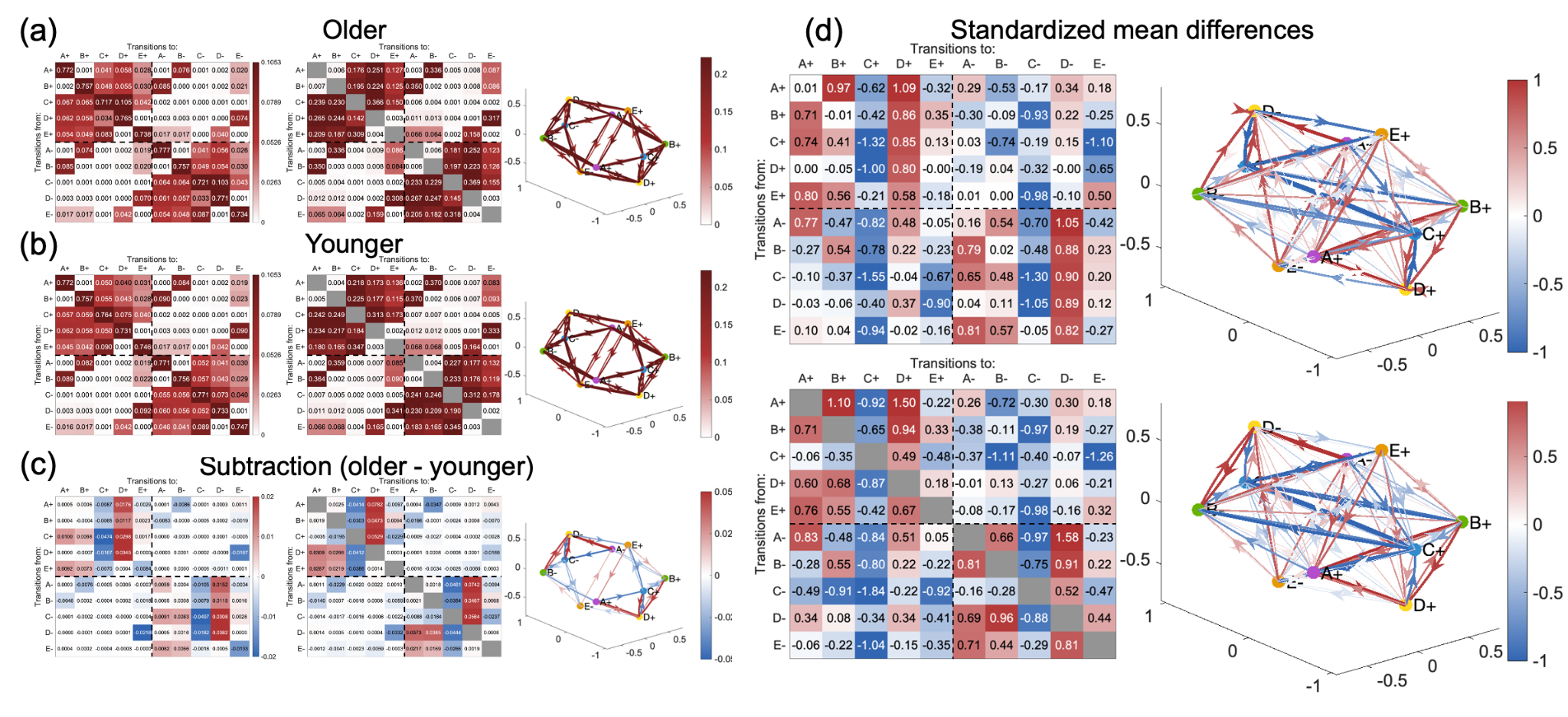

**Supplementary H: Results of confirming the transition component structure in Study 2**

To check whether the co-occurrence structure of the 100 transition pairs (i.e., transition components) was a robust feature, the eigenvectors obtained from the PCA in Study 1 and the rotation matrix obtained from subsequent Varimax rotation in Study 1 were applied to the data in Study 2. The rotated component loadings for each transition pair are shown in Table S2. Transition pairs were grouped as belonging to the component with the highest absolute value of rotated component loadings (highlighted in yellow in the table). Each transition component was named based on the content of the transition pairs to which it belonged. For convenience, each transition pair was labeled with positive states in upper case (e.g., A+ was denoted by “A”) and negative states in lower case (e.g., A- was denoted by “a”), in order of “from” and “to” states (e.g., the transition from A+ to A- was denoted by “Aa”).

**Table S2.** *Rotated component loadings of each transition pair in Study 2. Loading values were calculated by applying the eigenvectors and Varimax rotation matrix from Study 1.*

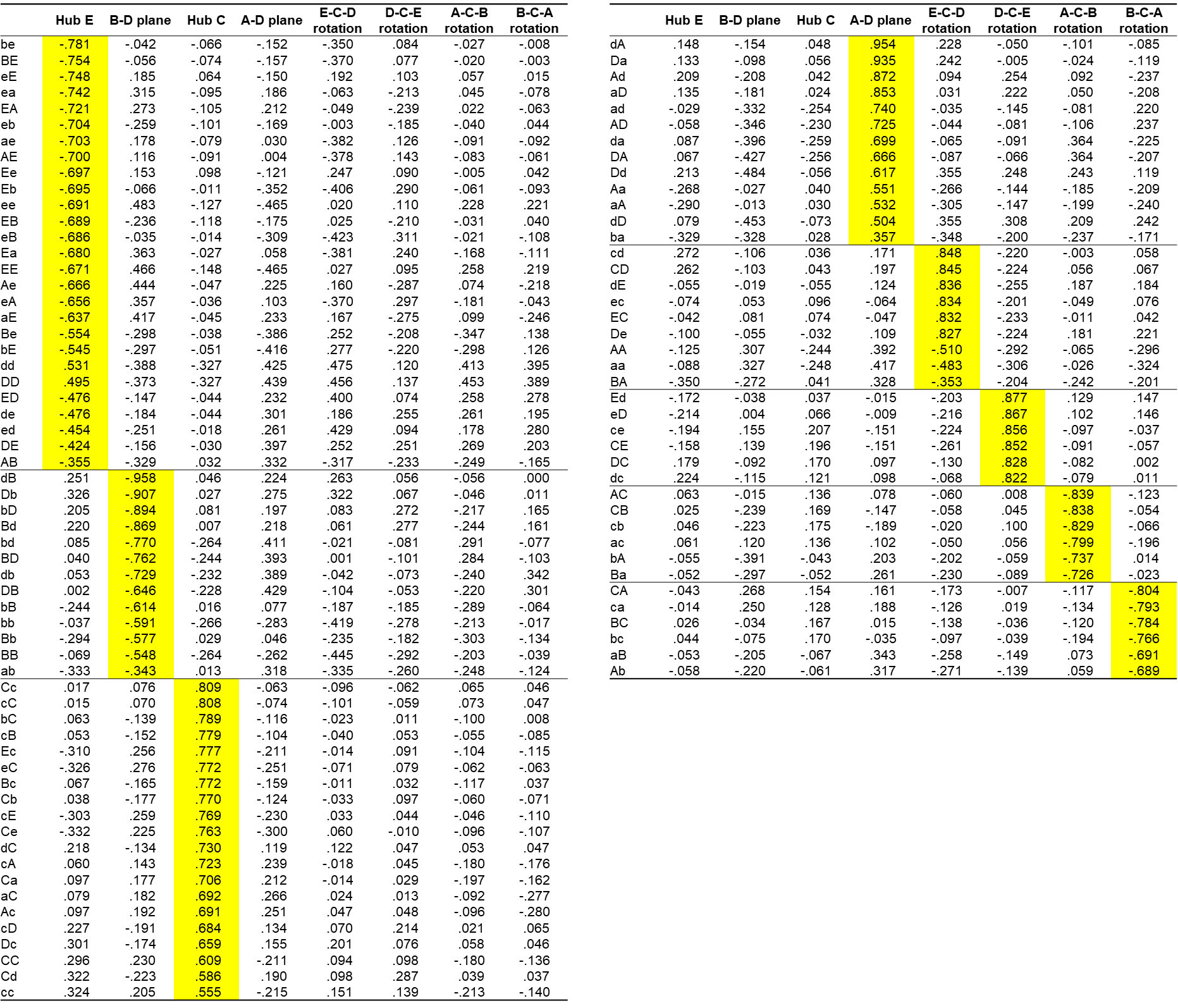
